# Limited predictability of amino acid substitutions in seasonal influenza viruses

**DOI:** 10.1101/2020.07.31.231100

**Authors:** Pierre Barrat-Charlaix, John Huddleston, Trevor Bedford, Richard A. Neher

## Abstract

Seasonal influenza viruses repeatedly infect humans in part because they rapidly change their antigenic properties and evade host immune responses, necessitating frequent updates of the vaccine composition. Accurate predictions of strains circulating in the future could therefore improve the vaccine match. Here, we studied the predictability of frequency dynamics and fixation of amino acid substitutions. Current frequency was the strongest predictor of eventual fixation, as expected in neutral evolution. Other properties, such as occurrence in previously characterized epitopes or high *Local Branching Index* (LBI) had little predictive power. Parallel evolution was found to be moderately predictive of fixation. While the LBI had little power to predict frequency dynamics, it was still successful at picking strains representative of future populations. The latter is due to a tendency of the LBI to be high for consensus-like sequences that are closer to the future than the average sequence. Simulations of models of adapting populations, in contrast, show clear signals of predictability. This indicates that the evolution of influenza HA and NA, while driven by strong selection pressure to change, is poorly described by common models of directional selection such as travelling fitness waves.

## INTRODUCTION

Seasonal influenza A viruses (IAV) infect about 10% of the global population every year, resulting in hundreds of thousands of deaths [1, 2]. Vaccination is the primary measure to reduce influenza morbidity. However, the surface proteins hemagglutinin (HA) and neuraminidase (NA) continuously accumulate mutations at a high rate, leading to frequent antigenic changes [2–5]. While a vaccine targeting a particular strain may be efficient for some time, antigenic drift will sooner or later render it obsolete. The World Health Organization (WHO) regularly updates influenza vaccine recommendations to best match the circulating strains. Since developing, manufacturing, and distributing the vaccine takes many months, forecasting the evolution of influenza is of essential interest to public health [6, 7].

The number of available high quality HA and NA sequences has increased rapidly over the last 20 years [8, 9] and virus evolution and dynamics can be now be tracked at high temporal and spatial resolution [10]. This wealth of data has given rise to an active field of predicting influenza virus evolution [6, 7]. These models predict the future population of influenza viruses by estimating strain fitness or proxies of fitness. Łuksza and Lässig [11], for example, train a fitness model to capture antigenic drift and protein stability on patterns of epitope and non-epitope mutations. Other approaches by Steinbrück et al. [12], Neher et al. [13] predict fitness by using hemagglutination inhibition (HI) data to determine possible antigenic drift of clades in the genealogy of the HA protein. Finally, Neher et al. [14] use branching patterns of HA phylogenies as a proxy for fitness. These branching patterns are summarized by the Local Branching Index (LBI), which was shown to be a proxy of relative fitness in mathematical models of rapidly adapting populations [14].

The underlying assumption of all these methods is that (i) differences in growth rate between strains can be estimated from sequence or antigenic data and (ii) that these growth rate differences persist for long enough to be predictive of future success. Specific positions in surface proteins are of particular interest in this context. The surface proteins are under a strong positive selection and change their amino acid sequence much more rapidly than other IAV proteins or than expected under neutral evolution [4, 15]. Epitope positions, i.e., positions targeted by human antibodies, are expected to change particularly often since viruses with altered epitopes can evade existing immune responses [3, 5, 16]. It therefore seems plausible that mutations at these positions have a tendency to increase fitness and a higher probability of fixation [15]. But one has to be careful to account for the fact that these positions are often ascertained post-hoc [3] and human immune responses are diverse with substantial inter-individual variation [17].

In this work, we use HA and NA sequences of A/H3N2 and A/H1N1pdm influenza from year 2000 to 2019 to perform a retrospective analysis of frequency trajectories of amino acid mutations. We quantify how rapidly mutations at different frequencies are lost or fixed and how rapidly they spread through the population. We further investigate whether any properties or statistics are predictive of whether a particular mutation fixes or not. To our surprise, we find that the predictability of these trajectories is very limited: The probability that a mutation fixes differs little from its current frequency, as would be expected if fixation happened purely by chance. This observation holds for many different categories of mutations, including mutations at epitope positions. This weak predictability is not attributable solely to clonal interference and genetic linkage, as simulation of models including even strong interference retain clear signatures of predictability. Consistent with these observations, we show that a simple predictor uninformed by fitness, the consensus sequence, performs as the well as the Local Branching Index (LBI), the growth measure based on the genealogy used in [14]. This suggests that although LBI has predictive power, the reason for its success may not be related to it approximating fitness of strains.

## RESULTS

The main underlying question asked in this work is the following: given a mutation *X* in the genome of influenza that we observe at a frequency *f* in the population at a given date, what can we say about the future of *X*? The trajectory of a mutation will depend on its own effect on fitness, the contribution of the genetic background on the same segment, and the effect of the remaining seven segments. Here, we investigate properties of broad categories of mutations effectively averaging over different genetic backgrounds to isolate the effects intrinsic to the mutation.

First, we ask whether we can quantitatively predict the frequency of *X* at future times *f* (*t*). In other words, having observed a mutation at frequencies (*f*_1_*, f*_2_*, …, f_n_*) at dates (*t*_1_*, t*_2_*, …, t_n_*), what can we say about its frequency at future dates (*t_n_*_+1_*, t_n_*_+2_*, …*)? A simpler, more qualitative question, is to ask whether *X* will fix in the population, will disappear, or whether the site will stay polymorphic.

We use amino-acid sequences of the HA and NA genes of A/H3N2 (since the year 2000) and A/H1N1pdm (since the year 2009) influenza available in GISAID [9] (see supplementary materials for an acknowledgment of all data contributors). This amounts to 44 976 HA and 36 300 NA sequences for A/H3N2 and 45 350 HA and 40 412 NA sequences for A/H1N1, with a minimum of 100 per year. These sequences are binned in non-overlapping intervals of one month. Each single-month time bin and the sequences that it contains represent a (noisy) snapshot of the influenza population at a given date. The number of sequences per time bin varies strongly both with year and according to the season, with earlier time bins containing around 10 sequences while more recent bins contain several hundreds (see figures S5 and S6 in SM for details).

The central quantities that we derived from this data are *frequency trajectories* of amino acids at each position in the sequences. If an amino acid *X_i_* is found at position *i* at a frequency between 5% and 95% in the population of a given time bin *t*, then the population is considered polymorphic at position *i* and at time *t*. This polymorphism is characterized by the frequency of *X_i_*, 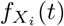, and also by frequencies of other amino acids at *i*. The series of values 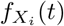 for contiguous time bins constitutes the frequency trajectory of *X_i_*. A trajectory is terminated if the corresponding frequency is measured above 95% (resp. below 5%) for two time bins in a row, in which case amino acid *X_i_* is considered as *fixed* (resp. *absent*) in the population. Otherwise, the trajectory is considered *active*. Examples of trajectories can be seen in figure S7 of the Supplement.

In the rest of this work, we will focus on frequency trajectories that are starting at a zero (low) frequency, *i.e. f* (*t* = 0) = 0. These represent new amino acid variants which were absent in the population at the time bin when the trajectory started and are currently rising in the population (see Methods). Such distinction in novel and ancestral variants is necessary to meaningfully interrogate predictability. Each rising trajectory of a new mutation implies the existence of another decreasing one at the same position, since frequencies of all amino acids at a given position must sum to one. If novel variants arise by selection, we expect to see a stronger signal of selection after conditioning on these novel variants. In classic models of population genetics, strongly advantageous variants undergo rapid selective *sweeps*, i.e., the rapid rise and fixation. The sweep of a mutation can be due to its own fitness effect, to the genetic background or to the effect of the seven other segments. By considering the ensemble of novel variants that are rising in frequency, we effectively average over backgrounds, obtaining a set of mutations that we expect to be beneficial on average. If such sweeps are common in the evolution of HA and NA, the restriction to trajectories that start at low frequency should thus enrich for mutations that are positively selected and on their way to fixation.

### Predicting future frequencies

Having observed the frequency trajectory *f* (*t*) of a mutation until a given date *t*_0_, how much can we say about the future values of *f* after *t*_0_? We consider the idealized case sketched in panel **A** of figure 1: given the trajectory of a *new* mutation, *i.e.* that started at a frequency of 0, and that we observe at frequency *f*_0_ at time *t*_0_, what is the probability *P*_Δ*t*_(*f*) of observing it at a value *f* at time *t*_0_ + Δ*t*?

**FIG. 1.**
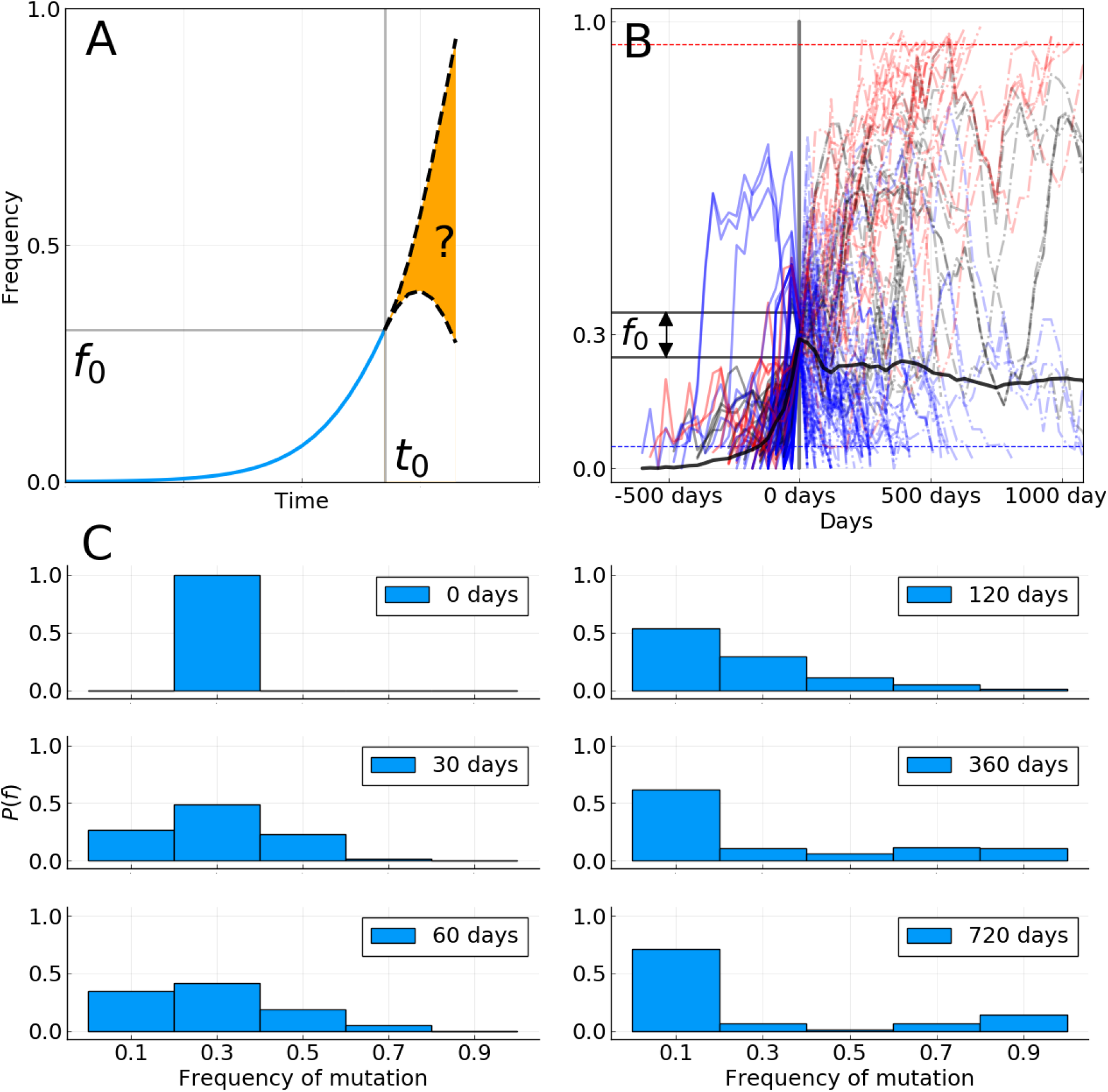
**A**: Sketch of the idea behind the short term prediction of frequency trajectories. Given a mutation that we have seen increasing in frequency and that we “catch” at frequency *f*_0_ at time *t*_0_, what can we say about the distribution of future frequencies *P*Δ*t*(*f| f*0)? **B**: All frequency trajectories of amino acid mutations in the A/H3N2 HA and NA genes that were absent in the past, are seen around *f*_0_ = 30% frequency at time *t*_0_ = 0, and are based on more than 10 sequences at each time point. Red curves represent mutations that will ultimately fix, blue the ones that will be lost, and black the ones for which we do not know the final status. Dashed horizontal lines (blue and red) represent loss and fixation thresholds. The thick black line is the average of all trajectories, counting those that fix (resp. disappear) as being at frequency 1 (resp. 0). Figure S8 shows equivalent figures for other values of *f* 0. **C**: Distribution of future frequencies *P*Δ*t*(*f| f*0) for the trajectories shown in panel **B** and for specific values of Δ*t*.

To answer this question retrospectively, we use all frequency trajectories extracted from HA and NA sequences that satisfy these conditions for a given *f*_0_. The number of trajectories is limited and the frequency estimates themselves are based on a finite sample and are hence imprecise. Therefore, we consider trajectories in an interval [*f*_0_ *− δf, f*_0_ + *δf*] with *δf* = 0.05.

For *f*_0_ = 0.3, we found 120 such trajectories in the case of A/H3N2 influenza, represented on the panel **B** of figure 1, where time is shifted such that *t*_0_ = 0. The same analysis was performed for A/H1N1pdm, with the 89 found trajectories displayed in figure S9. Some trajectories fall in the frequency bin around *f*_0_ while decreasing, even though they crossed that bin at an earlier time. This is due to the fact that some trajectories “skipped” the interval *f*_0_ in question on their initial rise due to sparse sampling. These trajectories are nevertheless rising in the sense that they start at frequency 0 for *t* → −∞. Removing them does not change results significantly.

Since rapid sequence evolution of influenza HA and NA mediates immune evasion, one could expect that a significant fraction of new amino acid mutations on rising trajectories in figure 1 are *adaptive*. We could thus expect that most of these trajectories continue to rise after reaching frequency *f*_0_, at least for some time. A fraction of those would then sweep through the population and fix.

To quantify the extent to which this preconception of sweeping adaptive mutations is true, we estimated the probability distribution *P*_Δ*t*_(*f| f*_0_) of finding a trajectory at frequency *f* after a time Δ*t* given that it was observed at *f*_0_ at time 0. The results for different Δ*t* are shown in figure 1C. Initially, *i.e.* at time *t*_0_ = 0, this distribution is by construction peaked around *f*_0_. If a large fraction of the trajectories keep increasing after this time, we should see the “mass” of *P*_Δ*t*_(*f| f*_0_) move to the right towards higher frequencies as time progresses.

However, future distributions for Δ*t >* 0 do not seem to follow a pattern compatible with selective sweeps. The thick black line in Figure 1B shows the average frequency of all trajectories. This average makes a sharp turn at *t* = 0 and is essentially flat for *t >* 0 in the case of A/H3N2, and slightly increasing for A/H1N1pdm (see supplement). Hence, the fact that this average rose for *t <* 0 gives little information for *t >* 0, and is due to the conditions by which these trajectories were selected. This shows that sweep-like trajectories rising steadily from frequency 0 to 1 are not common enough to dominate the average trajectory.

Consistent with the average, the frequency distribution of the selected trajectories broadens in time without a significant shift of the mean as time passes. After 60 days, the distribution is rather symmetrical around the initial *f*_0_ = 0.3 value, suggesting that the knowledge that the trajectories were rising is lost after two months. On a timescale of 60 to 120 days, the only possible prediction is that trajectories are likely to be found in a broad interval around the initial frequency *f*_0_. After one year the distribution becomes almost flat (excluding mutations that have disappeared or fixed), and the initial peak at *f*_0_ is not visible anymore. The only information remaining from the initial frequency is the fraction that fixed or was lost (see below). This behavior is expected in neutral models of evolution [18] but incompatible with a dynamic dominated by sweeps taking over the population.

While this observation does not rule out that signatures exist that predict future frequency dynamics, past dynamics alone is weakly informative.

### Prediction of fixation or loss

Instead of predicting future frequency, let’s consider the long-term goal of predicting the probability that a mutation fixes in the population. We first estimate the fraction of frequency trajectories that either fix in the population or are lost, as well as the time it takes for one or the other to happen. Panels **A** and **B** of figure 2 shows the fraction of frequency trajectories in HA and NA that either have fixed, were lost or remained active as a function of the time elapsed since they were first seen above 25% frequency. Most mutations are either lost or become fixed after 2-3 years, with very few trajectories remaining active after 5 years. This time scale of 2-3 years is consistent with the typical coalescence time observed in phylogenetic trees of A/H3N2 influenza [10, 19]. We also note that the fraction of lost trajectories increases sharply at small times with 40% of mutations observed above 25% frequency being lost within one year for A/H3N2, while it takes longer to fix a mutation in the whole population.

**FIG. 2.**
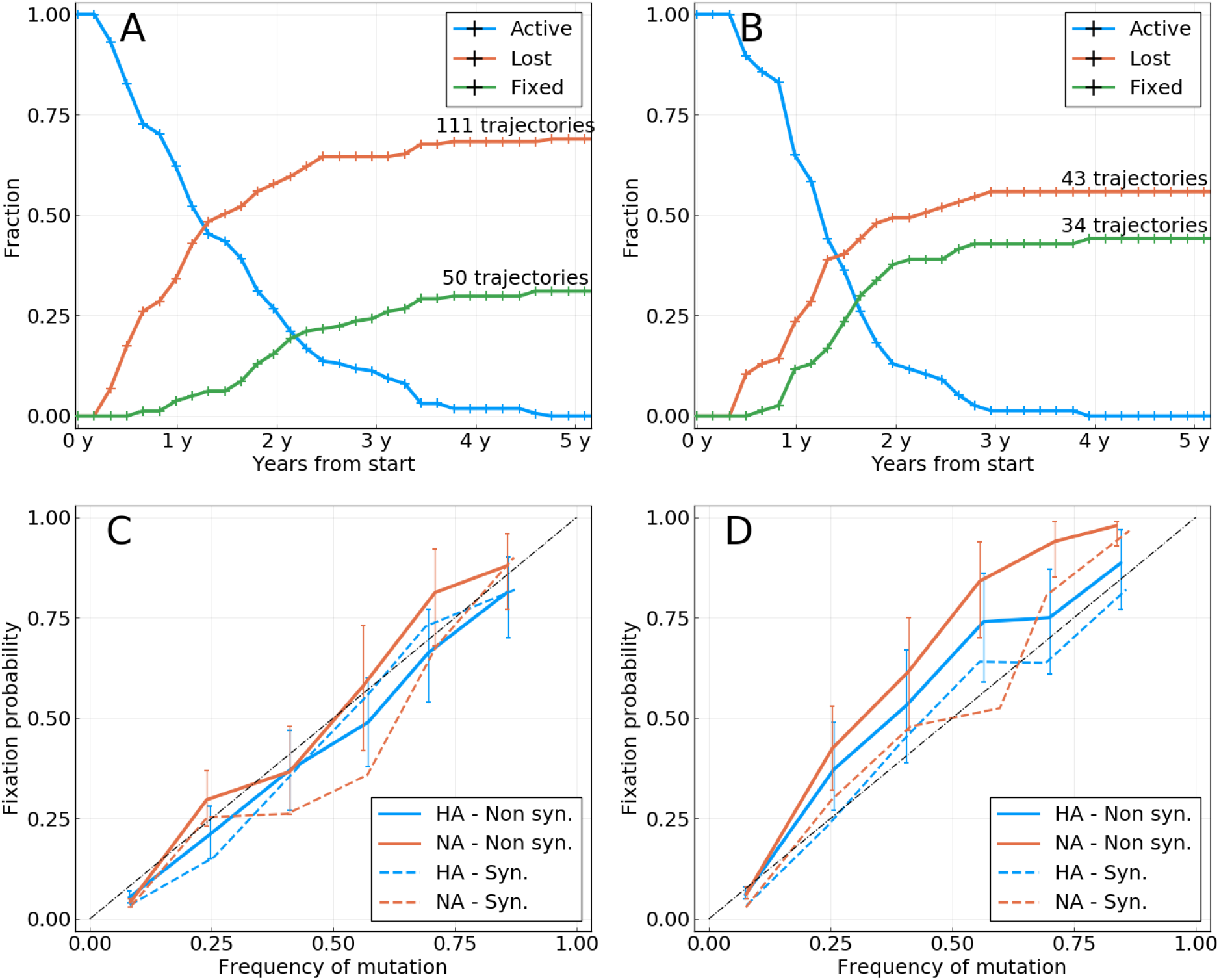
**A**: Activity of all rising frequency trajectories seen above 25% frequency for A/H3N2 HA and NA. **B**: Same as **A** for A/H1N1. **C**: Probability of fixation of a mutation (amino acid or synonymous) *P_fix_*(*f*) as a function of the frequency *f* at which it is measured, for A/H3N2 HA and NA. Only new mutations are considered, *i.e.* mutations that were absent in the past. The diagonal dashed line is the expectation from a neutrally evolving population. Colored dashed lines represent synonymous mutations. Colored solid lines represent amino acid mutations. Error bars represent a 95% confidence interval. **D**: Same as **C** for A/H1N1.

We then examined the probability of mutations to fix in the population as a function of the frequency at which they are seen. For different values of frequency *f*, we consider all trajectories that started at a null frequency and are seen in the interval [*f* − 7.5%*, f* + 7.5%] at any given time. The probability of a mutation fixing given that it is seen at frequency *f*, *P_fix_*(*f*), is then estimated by the fraction of those trajectories which terminate at a frequency larger than 95%, *i.e.* our fixation threshold. Panels **C** and **D** of figure 2 show *P_fix_*(*f*) as a function of *f* for NA and HA. For both proteins, the probability of fixation of a new mutation at frequency *f* is close to *f* itself, that is *P_fix_*(*f*) ≃ *f*. This result is exactly what is expected in a population evolving in the *absence* of selection. A mutation or trait appearing at frequency *f* is shared by *f · N* individuals, and the probability for one of them to become the ancestor of all the future population is *f · N/N* = *f*. Thus, the probability of this mutation or trait to fix in the population is equal to its current frequency, a case which we will refer to as the neutral expectation. Panel **C** of figure 2 indicates that mutations in the surface proteins of A/H3N2 influenza are in good agreement with the neutral expectation, while those in A/H1N1pdm show only small deviations from it. In both cases, the probability of fixation seems to be mainly dictated by the current frequency *f* at which the mutation is observed.

This dynamics is in apparent contradiction with evidence that influenza surface proteins are under strong selective pressure to evade human immune responses [4]. If strong selection was present, we would expect rising amino acid mutations to fix at a distinctively higher frequency than the one at which they are measured. In an extreme case where most trajectories would be clean sweeps, *P_fix_*(*f*) should be close to 1 for all but very small values of *f*.

Next, we searched for features of mutations that allow prediction of fixation beyond frequency by dividing frequencies into categories that deviate from the diagonal in panels **C** and **D** of figure 2. We first turn to the *Local Branching Index* (LBI), a quantity calculated for each node in a phylogenetic tree that indicates how dense the branching of the tree is around that node. LBI has previously been successfully used as a predictor of the future population of influenza [14], and was shown to be a proxy for fitness of leaves or ancestral nodes in mathematical models of evolution. Here, we define the LBI of a mutation at date *t* as the average LBI of strains that carry this mutation and that were sampled in the time bin corresponding to *t*. Panel A of figure 3 shows fixation probability for HA mutations with LBI in the top or bottom half of the distribution. Both groups have identical probability of fixation, suggesting that LBI carries very little information on the probability of fixation of a mutation.

**FIG. 3.**
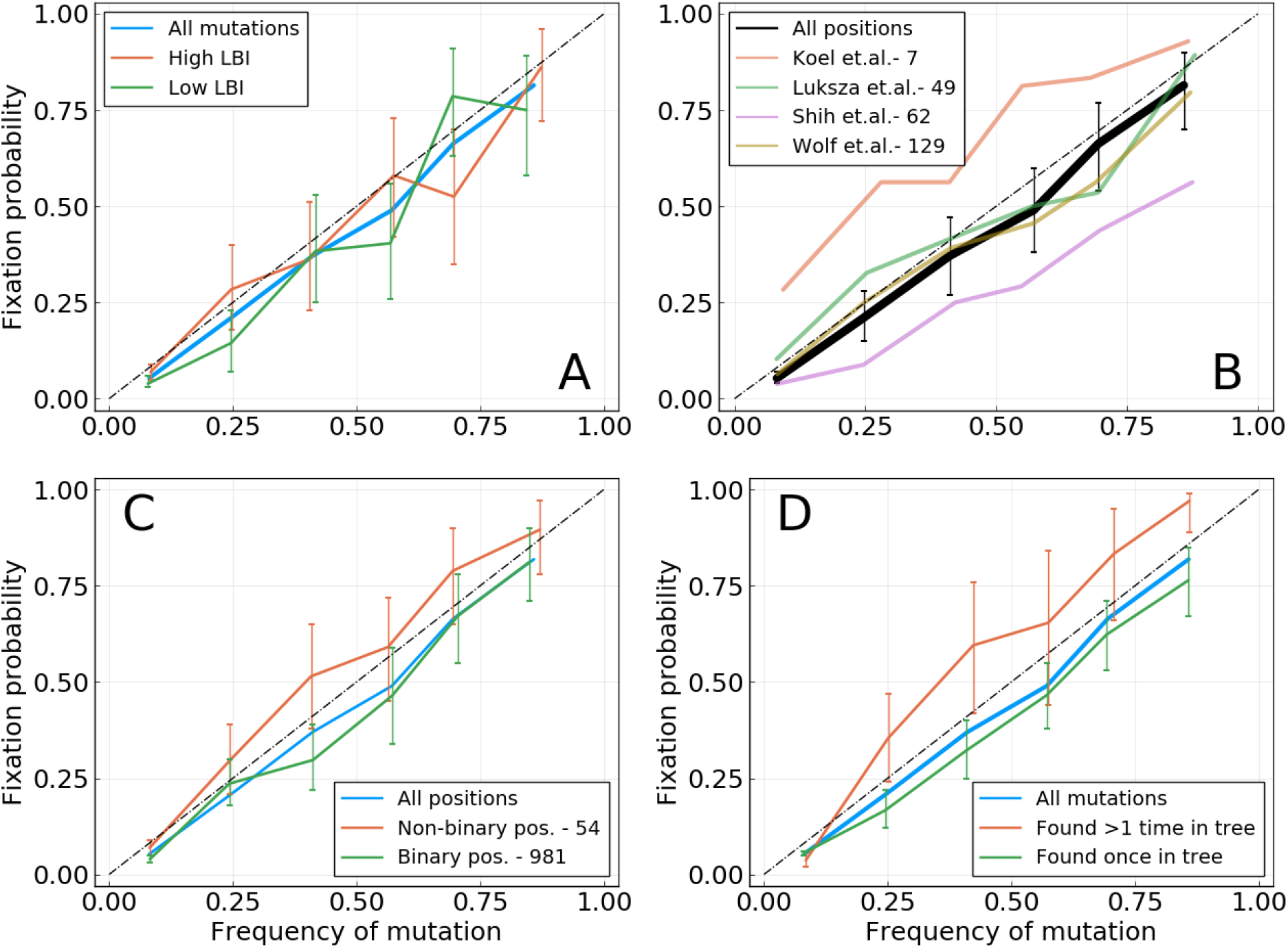
Fixation probability *P_fix_*(*f*) as a function of frequency, for A/H3N2 influenza. Figure S11 shows the same analysis for A/H1N1. **A**: HA mutations with higher or lower LBI values, based on their position with respect to the median LBI value. **B**: Different lists of epitope positions in the HA protein. The authors and the number of positions is indicated in the legend. **C**: HA and NA mutations for binary positions, *i.e.* positions for which we never see more than two amino acids in the same time bin. **D**: HA and NA mutations that appear once or more than once in the tree for a given time bin.

Next, we focused on previously reported antigenic sites in the A/H3N2 HA protein, referred to as *epitope* positions. Mutations at these position might mediate immune escape and are therefore likely under strong selection and show sweep-like behavior. We used four lists of relevant epitope positions from different sources comprising from 7 to 129 positions in the sequence of the HA1 protein [3, 5, 11, 16]. Panel Fig. 3**B** shows fixation probability as a function of frequency for the four lists of epitopes. Only mutations at the 7 epitope sites reported in [5] have higher chances of fixation than expected by chance. No clear difference is found for the lists by Łuksza and Lässig [11], Wolf et al. [16], while positions from Shih et al. [3] show lower chances of fixation. One should also note that many of these positions were determined post-hoc and might be enriched for positions that experienced rapid substitutions before the publication of the respective studies.

Two ways of categorising mutations, however, suggest some power to predict fixation. In panel Fig. 3**C**, we split trajectories into those occurring at binary positions where only two amino acid variants co-circulate and non-binary positions with more than two variants. Novel variants at non-binary positions, *i.e.* ones for which competition between three amino acids or more has occurred at least once, have a higher chance of fixation. In panel **D**, we separated mutations that appear more than once or only once in the reconstructed tree (see methods), and found that the former fix more often. Panels **C** and **D** show that it is possible to gain some information on the chance of fixation of a particular mutation, as was done in panel **B**. However, the predictive power remains small, with the “top” curves in panels **C&D** being very close to the diagonal.

We conduct the same analysis on A/H1N1pdm influenza, with results shown in figure S11. Results are qualitatively similar to those obtained for A/H3N2, with LBI giving little information and mutations at non-binary positions having a higher chance of fixation. Panel **D**differs between figures 3 and S11, with convergent evolution giving less information on fixation in the latter case. However, this could be due to the shorter time period over which A/H1N1pdm evolved, resulting in a shorter tree and less possibilities of convergent evolution. Indeed, error bars for mutations appearing multiple times in **D** of figure S11 are relatively large, indicating a lower amount of trajectories.

Since influenza is seasonal in temperate regions, geographic spread and persistence might be predictive of the success of mutations. We quantify geographic spread of a mutation by the entropy of its frequency distribution across regions (see methods) and its persistence by the age of the trajectory by the time it reaches frequency *f* Figures S12 and S13 show the fixation probabilities as a function of observed frequency for mutations classified according to these scores. The two scores also allow a quantitatively moderate distinction between mutations: for a given frequency *f*, mutations found in many regions or those that are older (in the sense that they have taken more time to reach frequency *f*) tend to fix more often than geographically localized mutations or more recent ones, but the effect is small. These two scores are in fact correlated, with older trajectories representing mutations that are more geographically spread, as can be seen in figure S14 of SM. However, it is important to note that sampling biases and heterogeneity across time and space (see supplementary figures S5 and S6) make answering such specific hypothesis challenging. Frequency of mutations might thus be amplified through different sampling biases, making the connection between geographic spread, seasonality and mutation frequency non-trivial to measure.

### Simulations of models of adaptation

The results shown in figures 2 and 3 are difficult to reconcile with the idea that seasonal influenza virus evolution is driven by rapid directed positive selection. One possible explanation for the weakly predictable behaviour of mutations (beyond their current frequency) might be tight genetic linkage inside each segment and strong competition between different adaptive mutations [15, 20]. We design a simple model of population evolution based on the `ffpopsim` simulation software to test this hypothesis [21]. The model represents a population of binary genomes of length *L* = 200 evolving in a fitness landscape that changes through time.

First, we use an additive fitness function, with sequence (*x*_1_ *… x_L_*) having a fitness ∑_*i*_ *h*_*i*_*x*_*i*_. This implies that for a given genome position *i*, the trait *x_i_* = −1 is favored if *h_i_ >* 0 whereas *x_i_* = − 1 is favored if *h_i_ <* 0. All *h_i_*’s have the same magnitude, and only their signs matter. Every Δ*t* generations, we randomly choose a position *i* and flip the sign of *h_i_*, effectively changing the fitness landscape. Individuals in the population now have the opportunity to make an adaptive mutation at site *i* giving them a fitness advantage 2 |*h*|. A “flip” at position *i* of the fitness landscape will decrease fitness of all individuals that carried the adapted variant at position *i* and increases the fitness of those that happened to carry a deleterious variant.

To increase competition between genomes, we designed a second model that includes epistasis. Once again, the baseline fitness of a genome is an additive function, this time with values of *h_i_* that do not change through time. In addition, we added a component that mimics immune selection. Every Δ*t* generation, we now introduce “antibodies” that target a specific sub-sequence of length *l* = 5, noted 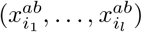. The positions (*i*_1_ *… i_l_*) are chosen at random, while the targeted sub-sequence is the dominant state at each position. Genomes that include the *exact* sub-sequence targeted by the antibody suffer a strong fitness penalty. However, a single mutation away from that sub-sequence removes this penalty completely, resulting in a fitness landscape with very strong epistasis. This has the effect of triggering a strong competition between adaptive mutations: for a given antibody, *l* = 5 possible mutations are now adaptive, but combinations of these mutations do not bring any fitness advantage.

Having simulated populations in these two fitness landscapes, we perform the same analysis of frequency trajectories as for the real influenza data. Figure S16 of the SM shows the *P_fix_*(*f*) as a function of *f* for the two models and for different values of the inverse rate of change Δ*t* of the fitness landscape. For all models, this curve deviates significantly from the diagonal. This is most evident for the case of a simple additive fitness landscape that changes rarely Δ*t* = 1000: rising mutations almost always fix in the population, with *P_fix_*(*f*) ≃ 1 for any *f* larger than a few percent. This is corroborated by visual inspection of the trajectories, which shows that evolution in this regime is driven by regular selective sweeps that take a typical time of ~400 generations. In other regimes, with smaller Δ*t* or with strong epistatic competition, *P_fix_*(*f*) is reduced and closer to the diagonal. However, it takes an extremely fast changing fitness landscape to push *P_fix_* close to the diagonal: with Δ*t* = 10, that is about 40 changes to the fitness landscape in the time it would take a selective sweep to go from 0% to fixation, *P_fix_*(*f*) differs from *f* in a way that is comparable to what is observed in A/H1N1pdm influenza.

These models are not meant to be accurate models of influenza viruses evolution. But figure S16 does show is that the patterns observed in influenza virus evolution are only reproduced by models of adapting populations when pushing clonal competition to extreme values. We conclude that the pattern in figure 2 may not be a straightforward manifestation of genetic linkage and clonal interference, but that some more intricate interplay of epidemiology, seasonality, human immunity and chance gives rise to the weakly predictable yet strongly selected evolutionary dynamics of IAVs.

### Why do predictions work?

The statistics of frequency trajectories seem to be in conflict with the notion that influenza evolution is predictable. Likewise, the LBI, a quantity that correlates with fitness in mathematical models and is used to predict future influenza populations [14], does not seem to contain any information on whether a specific mutation is going to fix or not, see figure 3. To resolve this conundrum, we first note that the criterion by which predictive power for influenza was measured in [14] was the distance between the strain with the highest LBI and the future population, not the ability of the LBI to predict dynamics. The distance was compared to the average distance between the present and future population, as well as the post-hoc optimal representative and the future.

To quantify the ability of the LBI and other measures to pick good representatives of the future, we construct a large tree of HA sequences with 100 sequences in nonoverlapping time bins of 4 months from year 2003 to 2019 (a total of 4402 as some 4 month intervals contain less than 100 sequences). Each time bin is considered as a snapshot of the A/H3N2 influenza population and we will refer to sequences in time bin *t* as the population of the *present*. From this present population, we predict *future* populations in time bin *t* + Δ*t*, using only sequences in time bin *t* and before.

To assess the ability of the LBI to pick a close representative of the future, we compute the LBI of each node of one time bin in the tree using only the leaves that belong to that time bin. The top panel in figure 4 shows the hamming distance of the strain with the highest LBI to future populations at different Δ*t* along with the same distance for a randomly chosen strain. The figure shows the distance averaged over all possible values of *t* for Δ*t* between 0 and 32 months, giving us an average efficiency of a predictor over 16 years of influenza evolution.

**FIG. 4.**
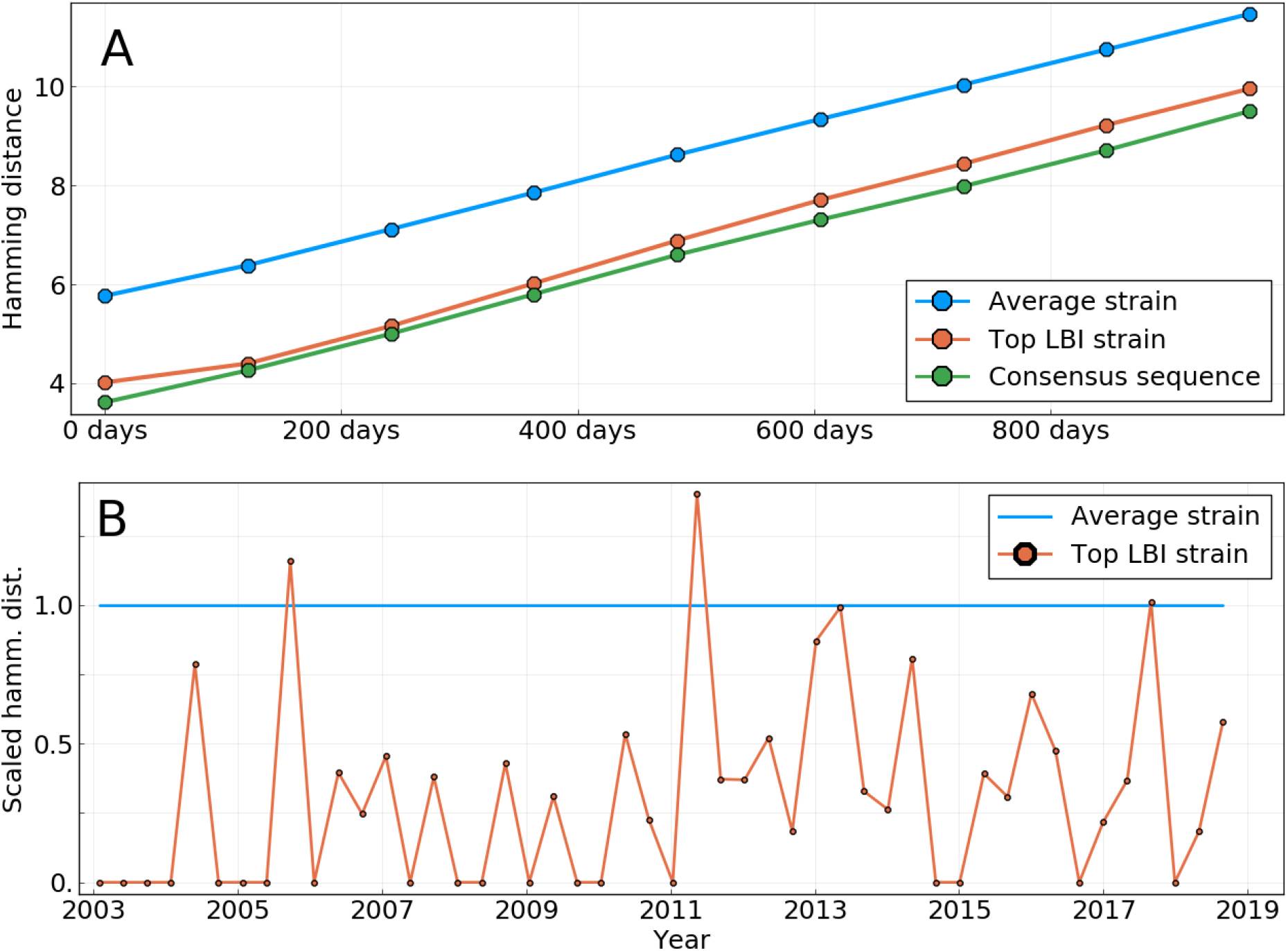
**A**: Average Hamming distance of the sequences of different predictors to HA sequences of future influenza populations, themselves averaged over all “present” populations from year 2003 to 2019. Predictors are: a randomly picked sequence in the present population; the sequence of the strain with the highest LBI in the present population; the consensus sequence of the present population. **B**: Scaled Hamming distance between the sequence of the top LBI strain and the consensus sequence for populations at different dates. The scaling is such that for each date, the Hamming distance between a strain from the population and the consensus is on average 1. The strain with the highest LBI is almost always closer to the consensus sequence than the average strain.

The strain with the highest LBI is consistently closer to the future than the average strain by about 1-2 amino acids, while the overall distance increases linearly due to the continuous evolution of the population. We hence reproduce previous results showing that the LBI picks closer than average representatives [14]. To investigate whether this apparent success is due to the ability of the LBI to predict fitness or not, we explored a different predictor: the amino acid consensus sequence of the present population (see Methods for a definition of the consensus sequence). The choice is motivated by the fact that it can be shown to be the best possible long term predictor for a neutrally evolving population in terms of Hamming distance (see SM section 1). Figure 4 shows that the consensus sequence is in fact a equally good or even slightly better representative of the future than the sequence with highest LBI (note that the consensus sequence does *not* necessarily exist in the population).

This near equivalence of the consensus and the strain with highest LBI can be explained as follows: The LBI tends to be high for nodes in a tree that are close to the root of a dense and large clade. A typical sample of influenza HA sequences fall into a small number of recognizable clades, and the strains with maximal LBI will often be close to the root of the largest of those clades. This root of the largest clade will often be close to the consensus of the whole population, explaining the similar distance patterns. To test that hypothesis, we measure the hamming distance from the sequence of the top LBI strain to the consensus sequence for populations of all time bins. Panel **B** of figure 4 shows these distances, scaled with respect to an average strain (details in caption). It clearly shows that the top-LBI strain and the consensus sequence are indeed quite similar: out of 48 time bins, only once is the sequence of the top-LBI strain farther away from the consensus than the average sequence is. Moreover, the sequence of the top-LBI strain *exactly* matches the consensus in 19 cases.

## DISCUSSION

Predicting the trajectory of a mutation requires (i) significant fitness difference between genomes carrying different variants at the site and (ii) a selection pressure that changes slowly over time. Under such conditions, it is expected that frequency trajectories will show a persistent behavior which would make them predictable for some time. However, we could find only limited evidence for such persistent behavior in the past 19 years of IAV evolution. This lead us to conclude that (i) influenza virus evolution is qualitatively different from models of rapidly adapting population (despite clear evidence for frequent positive selection), and (ii) previous methods to predict influenza evolution work primarily because they pick strains that represent the future well, not because they predict future dynamics.

The primary focus in this work was the investigation of frequency trajectories of new amino acid mutations. In the short term, we found that on average the direction of trajectory does not persist for longer than a few months. Indeed, the average trajectory in figure 1 takes a sharp turn when going from *t <* 0 to *t >* 0, instead of showing “inertia”. This suggests that selective sweeps are not representative of typical trajectories.

On a longer timescale, we investigated the probability that a novel mutation observed at frequency *f* fixes. In neutral models of evolution this probability equals *f*, while it should be higher or lower than *f* for mutations with a beneficial or deleterious effect on fitness, respectively. However, in the case of influenza, this probability differs little from *f*, making current frequency the best predictor for fixation. In figure 3, we split trajectories into groups for which we expected *P_fix_* to deviate from *f*. Many of these splits, such as high/low LBI or epitope/non-epitope positions, did not result in an increased predictability, while others gave limited information on fixation. Despite the lack of predictability of mutation frequency trajectories, influenza surface proteins show strong signatures of selection [4, 15].

Methods for predicting the future evolution of influenza either construct explicit fitness models [11, 22], use historical patterns of evolution [11, 23], phenotypic assays [13, 24], or dynamic or phylogenetic patterns [14, 25]. The goal of these methods is to pick strains that are good representatives of future populations and could serve as vaccine candidates [6].

The low power to predict frequency dynamics or fixation naturally triggers the question why the above methods have been found to work. Picking representatives of the future and predicting frequency dynamics are distinct objectives and success at the former (as compared to random picks) is not necessarily inconsistent with a lack of predictable dynamics. In fact, [22] reports that the rate at which the frequency of a strain changes is often a poor predictor – consistent with our observations here. But despite the fact that future frequencies are not predicted by the LBI, the strain with the highest LBI in the population is a better predictor of the future population than a randomly picked one. While the LBI was shown to be a correlate of relative fitness and be predictive of fixation in mathematical models of evolution [14], it does not seem to be predict influenza evolution because it measures fitness from genealogical structure. Instead, we believe it picks closer than average strains simply because it has the tendency to be maximal at the base of large and dense clades. These basal genotypes are closer to the future populations than the current tips of the tree and hence a better predictor on average. The consensus sequence of all present strains performs slightly but consistently better than picking the strain with the highest LBI. The consensus sequence is the best possible predictor for a neutrally evolving population, and does not attempt to model fitness in any way.

At the same time, influenza virus phylogenies show clear deviations from those expected from the neutral Kingman coalescent, similar to those expected under Bolthausen-Sznitman coalescent (BSC) processes that are generated by traveling wave models of rapid evolution [26, 27]. The correspondence between the BSC and traveling wave models comes from transient exponential amplification of fit strains before these fitness differences are wiped out by further mutation. This exponential amplification generates long-tailed effective offspring distributions which in turn can leads to genealogies described by the BSC [26, 28]. Many processes other than selection, including seasonality and spatio-temporal heterogeneity, can generate effective long tailed offspring distributions even in absence of bona-fide fitness differences, which might explain ladder-like non-Kingman phylogenetic trees.

A recent preprint proposed that influenza virus evolution is primarily limited by an asynchrony between population level selection and generation of new variants within infected hosts [29]. Along these lines, it is possible that the A/H3N2 population readily responds once population level selection is high enough by giving rise to essentially equivalent variants. Furthermore, selection might cause the rapid rise of a novel variant to macroscopic frequencies (observable in a global sample) but its benefit rapidly “expires” because competing variants catch up and/or it mediates immune escape only to a small fraction of the population. These considerations might explain the disconnect between models of rapid adaptation and the frequency dynamics observed in influenza virus populations.

## METHODS

### Data and code availability

The sequences used are obtained from the GISAID database [9]. Strain names and accession numbers are given as tables in two supplementary files. Outliers strains listed at https://github.com/PierreBarrat/FluPredictibility/src/config were removed.

The code used to generate the figures presented here is available at https://github.com/PierreBarrat/FluPredictibility.

### Frequency trajectories

For a set of sequences in a given time bin, we compute frequencies of amino acids at each position by simple counting. We make the choice of not applying any smoothing method in an attempt to be as close to the data and “model-less” as possible. This is especially important for the short term prediction of frequency trajectories, as estimations of the “persistence time” of a trajectory might be biased by a smoothing method.

We compute frequency trajectories based on the frequencies of amino acids. A trajectory begins at time *t* if an amino acid is seen under the lower frequency threshold of 5% (resp. above the higher threshold of 95%) for the two time bins preceding *t*, and above this lower threshold (resp. below the higher threshold) for time bin *t*. It ends in the reciprocal situation, that is when the frequency is measured below the lower threshold (resp. above the higher threshold) for two time bins in a row.

In order to avoid estimates of frequencies that are too noisy, we only keep trajectories that are based on a population of at least 10 sequences for *each* time bin. As said in the Results section, we also restrict the analysis to trajectories that begin at a 0 frequency, in part to avoid double counting. We find a total of 460 such trajectories. However, only 106 reach a frequency of 20%, on which figure 2 is based for instance.

Note that the fact that we use samples of relatively small sizes – at least for some time bins – leads to biases in the estimation of frequencies. We show in Supplementary Material that these biases are generally small and do not induce any qualitative changes to results presented here.

### Local Branching Index

LBI was introduced in [14] as an approximation of fitness in populations evolving under persistent selective pressure that is fully based on a phylogenetic tree. It relies on the intuition that the tree below high-fitness individuals will show dense branching events, whereas absence of branching is a sign of low-fitness individuals. Quantitatively, the LBI *λ_i_*(*τ*) of a node *i* is the integral of all of the tree’s branch length around *i*, with an exponentially decreasing weight *e^−t/τ^* with *t* being the branch length. When considering a time binned population, the LBI is computed once for each time bin by considering only the leaves of the tree that belong to the time bin. This means that only branches that ultimately lead to a leaf that belongs to the time bin are considered in the integration.

*τ* is the time scale for which the tree is informative of the fitness of a particular node. Here, we use a value of *τ* equal to a tenth of *T_C_* ≃ 6 years, the coalescence time for influenza A/H3N2 strains, converted to units of tree branch length through the average nucleotide substitution rate (≃4 · 10^*−*3^ substitutions per site per year for HA). We have observed that given our method to predict the future from present populations corresponding to time bins of 4 months, changing the value of *τ* has little effect on the pick of the top LBI strain. By retrospectively optimizing its value, it is possible to reduce the average distance to the population 2 years ahead by ~0.25 amino acids on average, making the LBI method almost as good as the consensus on figure 4.

### Measuring the geographical spread of a mutation

For a mutation *X* we define its regional distribution using the numbers *n_r_*(*X*) that represent the number of sequences sampled in region *r* that carry *X*. Regional weights are then defined as

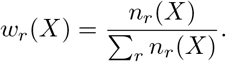

We can then measure the geographical spread *G*(*X*) of *X* by using the Shannon entropy of the probability distribution *w_r_*(*X*):

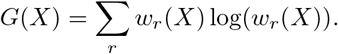

*G*(*X*) is a positive quantity that is larger when *X* is equally present in many regions, and equal to zero when *X* is concentrated in only one region.

Region used are the ones defined in the `Nextstrain` tool [30]. Those are North America, South America, Europe, China, Oceania, Southeast Asia, Japan & Korea, South Asia, West Asia, and Africa.

### Assigning a fitness to trajectories

#### Consensus sequence

Given a set of *N* sequences (*σ*^1^*, …, σ^N^*) based on an alphabet 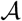 (*e.g.* 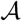 has 20 elements for amino acids, 4 for nucleotides), we can define a *profile* distribution *p_i_*(*a*) by the following expression:

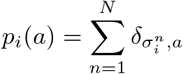

where *i* is a position in the sequence, 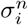 the character appearing at position *i* in sequence *σ*^n^, *a* a character of the alphabet and *δ* the Kronecker delta. The profile *p_i_*(*a*) simply represents the fraction of sequences which have character *a* at position *i*.

We then simply define the consensus sequence *σ^cons^* such that

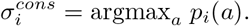

In other words, the consensus sequence is the one that has the dominant character of the initial set of sequences at each position.

### Earth Mover’s Distance

In order to measure the distance of several predictor sequences to the future population, we rely on the *Earth Mover’s Distance* (EMD), a metric commonly applied in machine learning to compare collections of pixels or words [31, 32]. Here, we apply it to compute the distance between the sequences of two populations, noted as 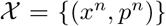 and 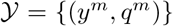 with *n* ∈ {1 *… N*} and *m* ∈ {1 *… M*}. In this notation, *x^n^* and *y^m^* are sequences, and *p^n^* and *q^m^* are the frequencies at which these sequences are found in their respective populations. For convenience, we also define *d_mn_* = *H*(*x^n^, y^m^*) as the Hamming distance between pairs of sequences in the two populations.

We now introduce the following functional

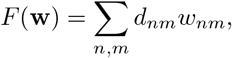

with **w**= *w_nm_* being a matrix of positive weights. The EMD between the two populations 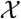 and 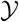 is now defined as the minimum value of function *F* under the conditions

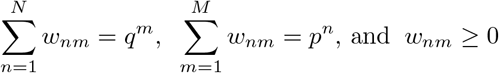

Intuitively, the weight *w_nm_* tells us how much of sequence *x^n^* is “moved” to sequence *y^m^*. The functional *F* sums all of these moves and attributes them a cost equal to the Hamming distance *d_nm_*. The conditions on weights in **w**ensure that all the weight *p^n^* of *x^n^* is “moved” to elements in 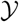 and vice versa.

The minimization is easily performed by standard linear optimization libraries. Here, we use the Julia library JuMP [33].

## Supporting information

Strains A/H3N2 NA

Strains A/H3N2 HA

Strains A/H1N1pdm NA

Strains A/H1N1pdm HA

## SUPPLEMENTARY MATERIAL

### 1. Consensus sequence as a predictor for neutrally evolving populations

We consider the case of a neutrally evolving and structure-less population, such as the one in the Wright-Fisher model of evolution [34]. At an initial time *t* = 0, the population consists of *N* individuals with genomes (*σ*^1^ *… σ^N^*) of length *L* (not necessarily distinct).

We make two hypotheses about this population. We first suppose that *no* mutations occur during the evolution of this population. This may seem surprising and is of course not true in the case of influenza. This assumption is however in line with the fact that the object of this work is to predict the outcome of *already existing* mutations in the influenza population. The prediction of mutations that we have not yet seen is not in its scope. Thus, assuming that no new mutations take place can be seen as a simple way to model the fact that we have no information about such events. The second assumption is that the population evolves in a completely neutral way, meaning that the average number of descendants of each genome *σ^n^* is the same. Let us now consider the population after it has evolved for a long time *t ≫ T* where *T* is the typical coalescence time (for the Wright-Fisher model, *T* = 2*N*). At this point, all individuals in the future population will descend from a unique individual *n*_0_ in the *t* = 0 population. Our two hypotheses now allow us to make two statements. First, since no new mutations are allowed, the population at *t ≫ T* will be clonal, with all individuals having genome *σ*^*n*0^. Second, since the evolution is neutral and does not favour any genome in particular, the probability that *σ*^*n*0^ is equal to a given genome *σ* is 1*/N*. In other words, the probability that a genome at *t* = 0 ultimately becomes the ancestor of all the future population is equal to its frequency in the *t* = 0 population.

We now try to find the genome *σ* that best predicts the future population on the long run, that is for *t ≫ T*. Here, we take best to mean that the predictor minimizes *H*(*σ, σ*^*n*0^) where *H* is the Hamming distance defined by

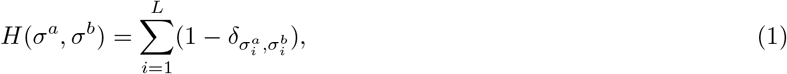

with *σ_i_* being the character appearing at position *i* of genome *σ* and *δ* the Kronecker delta. Since we do not know *n*_0_, we have to average over all its possible values. *σ* must thus minimize the following quantity:

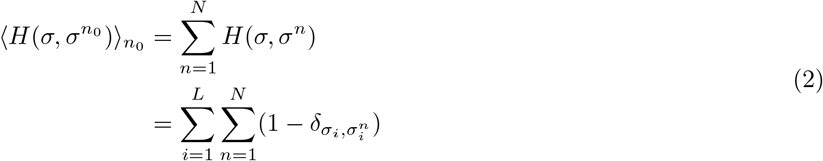

by using the definition of the Hamming distance. We now assume that characters at each positions of the genomes can be indexed by an integer *a* running from 1 to *q*. For instance, if these were amino acid sequences, we could index the 20 amino acids by *a* running from 1 to *q* = 20. We rewrite the Kronecker delta in the previous expression using this indexation:

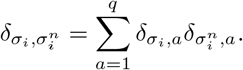

We also introduce the *profile* frequencies *p_i_*(*a*) of the population at time *t* = 0:

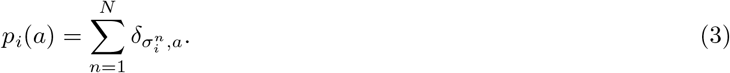

*p_i_*(*a*) represents the frequency at which character *a* appears at position *i* in genomes of the initial population. Equation 2 now becomes

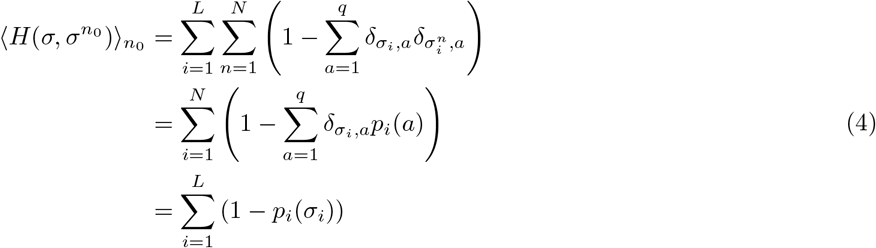

This means that the genome *σ* = (*σ*_1_ *… σ_L_*) which best predicts the future population according to our definition is the one that minimizes the quantity (1 − *p_i_*(*σ_i_*)) for all positions *i*. This obviously implies that each *σ_i_* must be chosen as to maximize *p_i_*(*a*), that is *σ_i_* must be the character that appears the most frequently at position *i*. Thus, *σ* must be the *consensus* sequence of the initial population.

### Predictor based on the local LBI maxima

In figure 15, we use several sequences as a predictor of the future population. Distance between two sets of sequences, *i.e.* the predictor sequences and the ones of the future population, is defined as the Earth Mover’s Distance (EMD). Here, we show that for a population evolving under the same hypotheses as in section 1, the best *multiple* sequence long term predictor is again the consensus sequence with weight 1.

Let the predictor be a set of weighted sequences (*s^α^, q_α_*). We again use the fact that in the long term, a unique sequence *σ*^*n*0^ from the present will be the ancestor of the entire population. We want to compute the EMD from the predictor to *σ*^*n*0^, that is the EMD between the sets 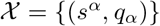 and 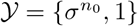. Applying the definition of the Methods section, it follows that the weights **w**are in this case equal to the *q_α_*s. By averaging over all values of *n*_0_, we now obtain

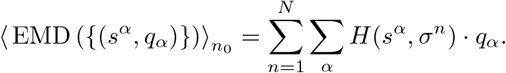

By the same calculation procedure as in the previous section, this expression simplifies to

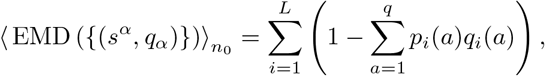

where the profile of the present population *p_i_*(*a*) has already been defined, and *q_i_*(*a*) stands for the profile of the predictor, that is

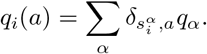

To minimize this distance, we find a profile *q*_i_ (*a*) that maximizes the quantity 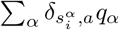 for each position *i*. It is clear that this is done by assigning a value *q_i_*(*a*) = 1 if *a* maximizes *p_i_*(*a*), and *q_i_*(*a*) = 0 otherwise. Thus, the profile of the predictor must be that of the consensus sequence, which is only possible if the predictor becomes *{σ^cons^,* 1*}*.

### 3. Biases in frequency estimations

The frequency of mutations in a given time-bin is simply performed by computing their frequency in sequences sampled in that time bin. This leads to potential biases in estimating frequencies, that arise for two reasons:

i. A mutation present at frequency *p* in the population might be observed at another frequency *f* ≠ *p* if *f* is estimated using a sub-sample of the population.
ii. For a neutrally evolving population, the distribution of frequencies of alleles is of the form *P* (*p*) ∝ 1*/p*. This means that the amount of alleles at frequency *p* is lower when *p* is higher.

**Figure S1.**
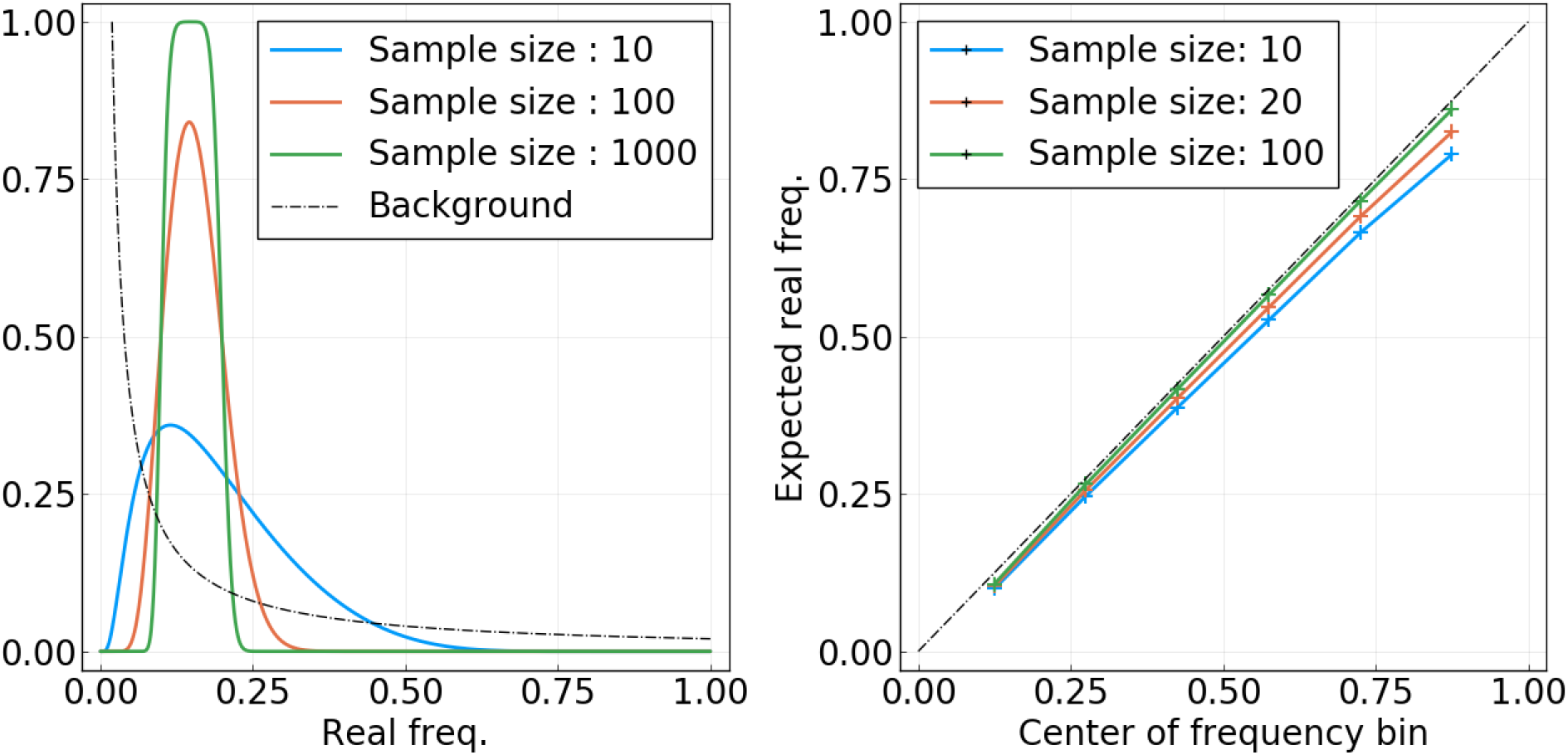
**Left**: For a mutation present at frequency *p* in the population, probability of being observed in the frequency bin [0.1, 0.2] as a function of *p* and for different sample sizes *n*. The dashed black line sketches the (non-normalized) background distribution *P_b_*(*p*). **Right**: Expected “real” average frequency of mutations found in frequency bin [*f*_1_*, f*_2_] as a function of the centre of the bin (*f*1 + *f*2)*/*2, for different sample sizes.

To illustrate *(i)*, let us compute the probability that a mutation present at “real” frequency *p* in the population is found to be in a given frequency bin [*f*_1_*, f*_2_] when *p* is estimated from a sample of size *n*. The sample consists of *n* observations {*x_i_*} with 1 ≤ *i ≤ n*, with *x_i_* = 1 if sequence sequence *i* of the sample bears the mutation, and *x_i_* = 0 if not. If *n* is small with regard to the total population size, we can consider the *x_i_* as random variables with a binomial distribution, meaning that *P* (*x_i_* = 1) = *p* and *P* (*x_i_* = 0) = 1 − *p*. The empirical frequency *f* is then estimated by taking the average of the *x_i_* variables, that is *f* = (*x*_1_ + *…* + *x_n_*)*/n*. If those are independently sampled and *n* is large enough, the probability of measuring value *f* is given by the Central Limit Theorem:

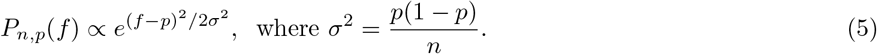

To compute the probability that this mutation is found in a given frequency bin [*f*_1_*, f*_2_], we integrate this distribution:

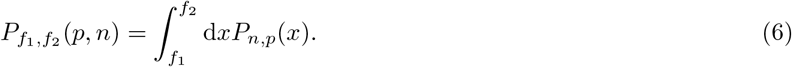

Function *P_f_*_1 *,f*2_ (*p, n*) is shown as a function of *p* for a fixed interval and for different values of *n* in the first panel of figure S1. Note the asymmetry of it: the variance of a binomial distribution of parameter *p* is small when *p* is close to 0 or 1, and goes through a maximum at *p* = 0.5. For this reason, mutations present at frequency *p* close to 0.5 have a higher probability of being observed in other frequency bins. On the contrary, this is unlikely for very rare or very frequent mutations.

We now try to estimate biases in frequency estimation due this phenomenon. Given a set of mutations that have been measured in frequency bin [*f*_1_*, f*_2_], what is the average *real* frequency of these mutations? To compute this, we need to sum *P_f_*_1 *,f*2_ (*p, n*) over all possible real frequencies *p*, giving us the amount of mutations that are observed in interval [*f*_1_*, f*_2_], and weigh this sum by the frequency value *p* as well as by the background distribution of frequencies *P_b_*(*p*) ∝ 1*/p*. This last quantity represents the expected amount of mutations that are present at frequency *p* in the population. Note that there is no divergence problem as the smallest non zero frequency is 1*/N*, where *N* is the population size. This leads us to the following expression for the average of “real” frequencies:

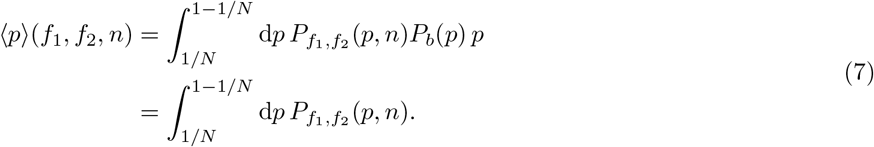

We have not made normalization explicit in these equations. It is simply achieved by dividing the above expression by ∫d*p P_f_*_1 *,f*2_ (*p, n*)*P_b_*(*p*).

In the second panel of figure S1, ⟨*p*⟩ (*f*_1_*, f*_2_*, n*) is plotted as a function of the centre of the interval [*f*_1_*, f*_2_] and for different values of *n*. For sample sizes *n >* 100, the biases due to this effect are almost non existent. For smaller samples, for instance *n* = 10, they are small but non negligible. However, we argue that this is not a significant problem with respect to the main results presented in this article. First, figure S6 shows that sample sizes of the order of *n* = 10 are only the case for a few months in the period going from year 2000 to 2018. From 2010 and onwards, more than a hundred sequences are available per month for most months. Secondly, even if most samples were in the *n* = 10 case, deviations shown in figure S1 are small enough that results shown in figures 2 and 3 would be *qualitatively* unchanged. Note that using the centre of the interval as a reference in figure S1, *i.e.* (*f*_1_ + *f*_2_)*/*2, would be correct in the case of a very large *n* and a flat background distribution *P_b_*(*p*). For figures 2 and 3 of the main text however, the average frequency of mutations found in an interval [*f*_1_*, f*_2_] is computed by taking the average of the observed frequencies, and not the centre of the interval. This partially takes into account biases considered here, as the background distribution *P_b_*(*p*) is then accounted for, even though it is equivalent to assuming infinite sample sizes.

**Figure S2.**
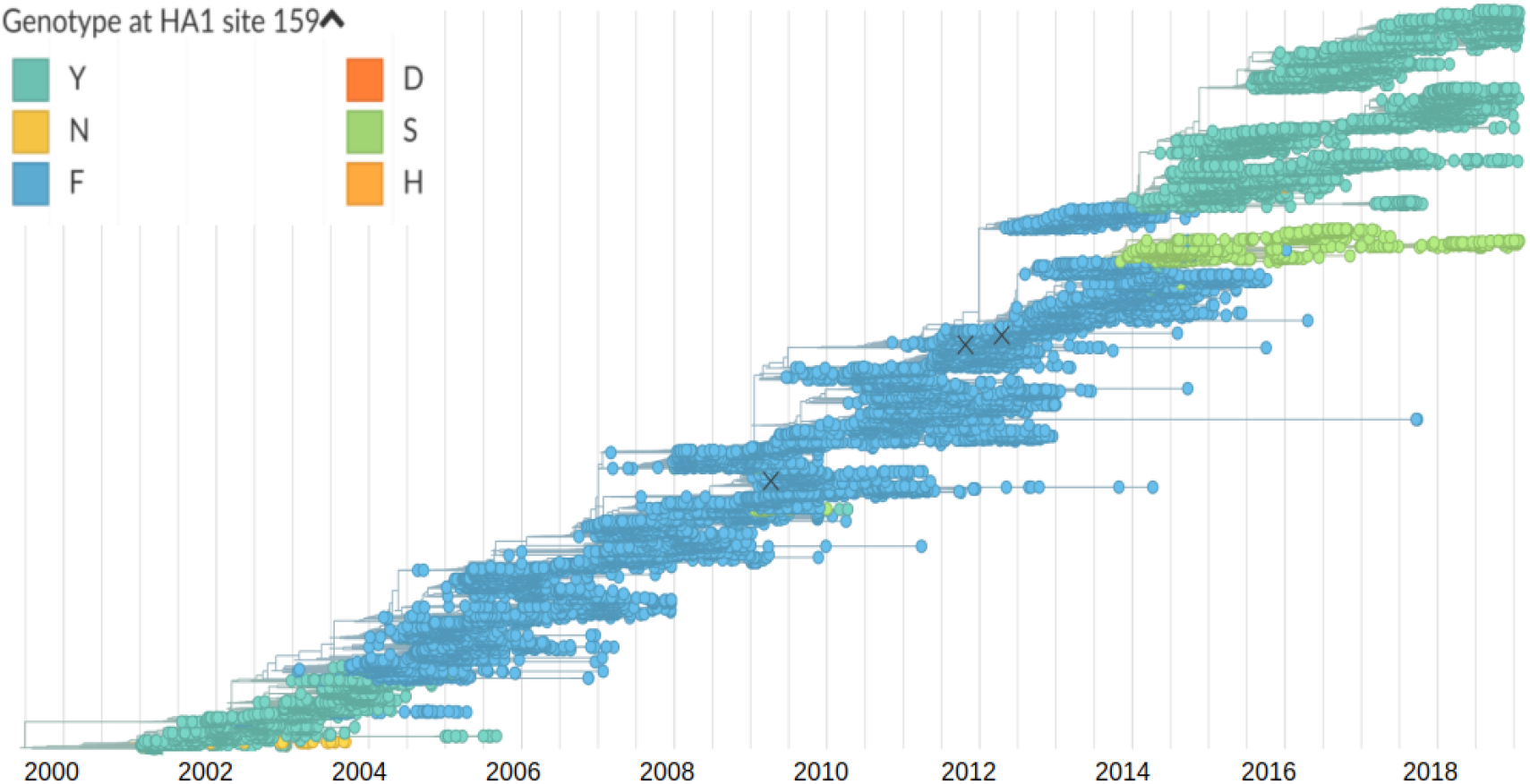
Tree used for this study, based on a random selection of 100 strains per month from year 2002 to 2018. Nodes and branches are colored according to the amino acid found at position HA1:159. The HA1 159S mutation is visible as a thin but long light-greened color branch, coalescing with the “trunk” around year 2013.

The analysis of the main text is in a large part based on the probability of fixation of mutations. The motivation underlying this choice is the relatively short coalescence time of the A/H3N2 influenza population, typically around three years. This can be seen in figure 2 of the main text, which shows the typical lifetime of frequency trajectories, ending in fixation or loss after at most 3 years in most cases. The tree in figure S2 is another illustration of this: for the most part of it, a “trunk” is clearly identifiable, and lineages that depart from it have a relatively short lifetime. This is no longer the case since the year ~2013: two clades have been competing since then, with no definite way to identify a trunk in the tree. The clade defined by the HA1 159S mutation, colored in light green on figure S2, is one of these two competing lineages. Because of this particular situation, the number of mutations fixating in the population is strongly reduced, as a mutation must appear in both clades to reach a frequency of 1. This is a potential flaw in our analysis, which concentrates on mutations fixating.

For this reason, we decided to re-run our analysis after having cut off the HA1 159S clade. In other words, we remove from the set of sequences those that carry the HA1 159S mutation. Results are shown in figures, equivalent to figures 2 and 3 of the main text. It is clear that qualitative results are left unchanged when this competing clade is removed. This can be surprising, as almost no complete fixation of an amino acid mutation has occurred since 2013. Cutting off the HA1 159S branch should thus result in many new fixations, changing the analysis. The reason for the similarity of results can be explained: fixation (resp. loss) of a mutation are defined here as the frequency of this mutation being measured above 95% (resp. 5%) frequency for two months in a row. As the HA1 159S clade is rather sparsely populated, it reaches frequencies lower than 5% two times (in 2015 and 2017), allowing mutations in the competing clade to “fix” as defined here. Thus, removing strains carrying HA1 159S does not introduce a significant amount of “new” fixation events.

**Figure S3.**
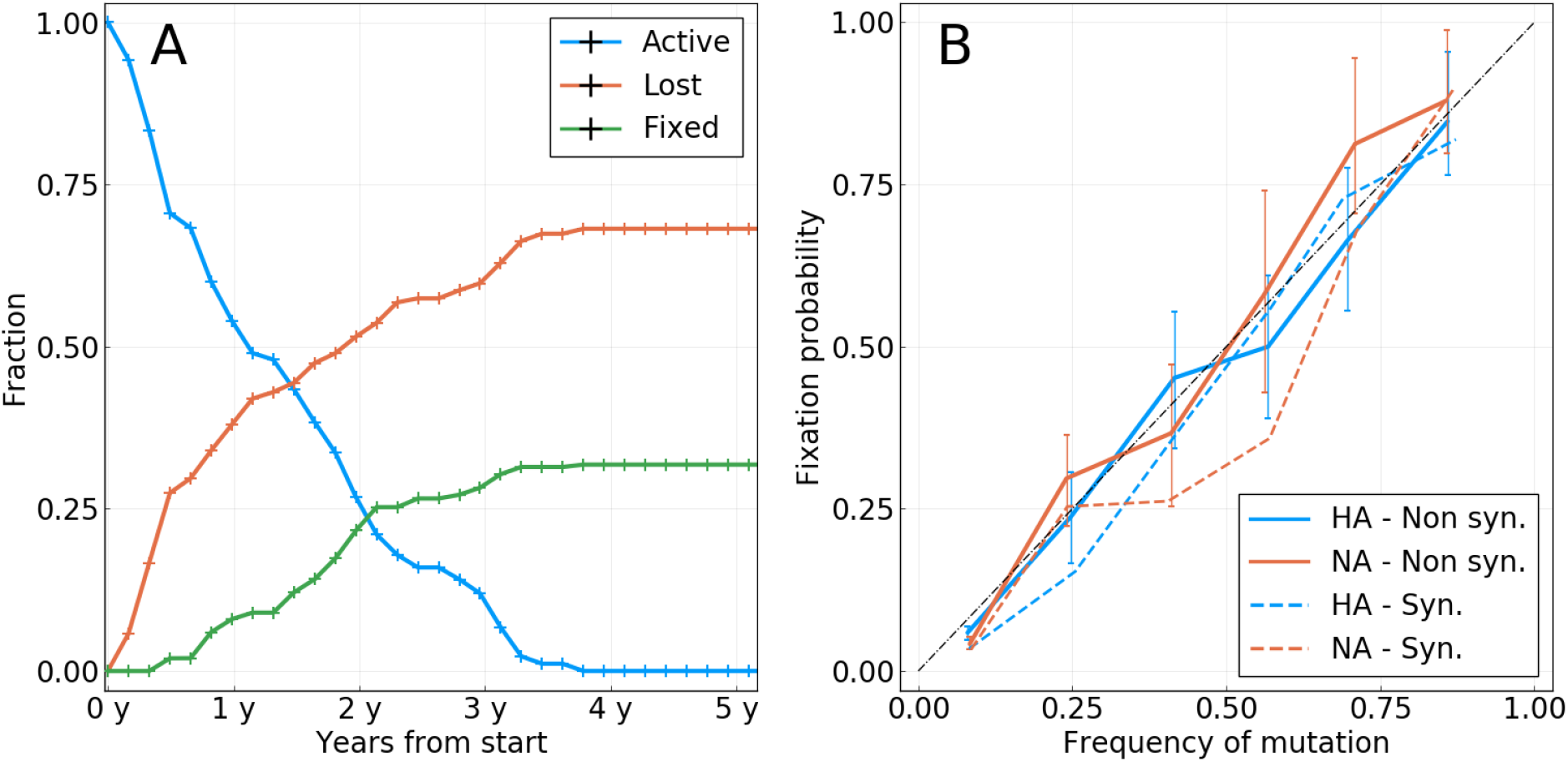
Equivalent to figure 2 of the main text, but with strains carrying the HA1 159S mutation removed.

**Figure S4.**
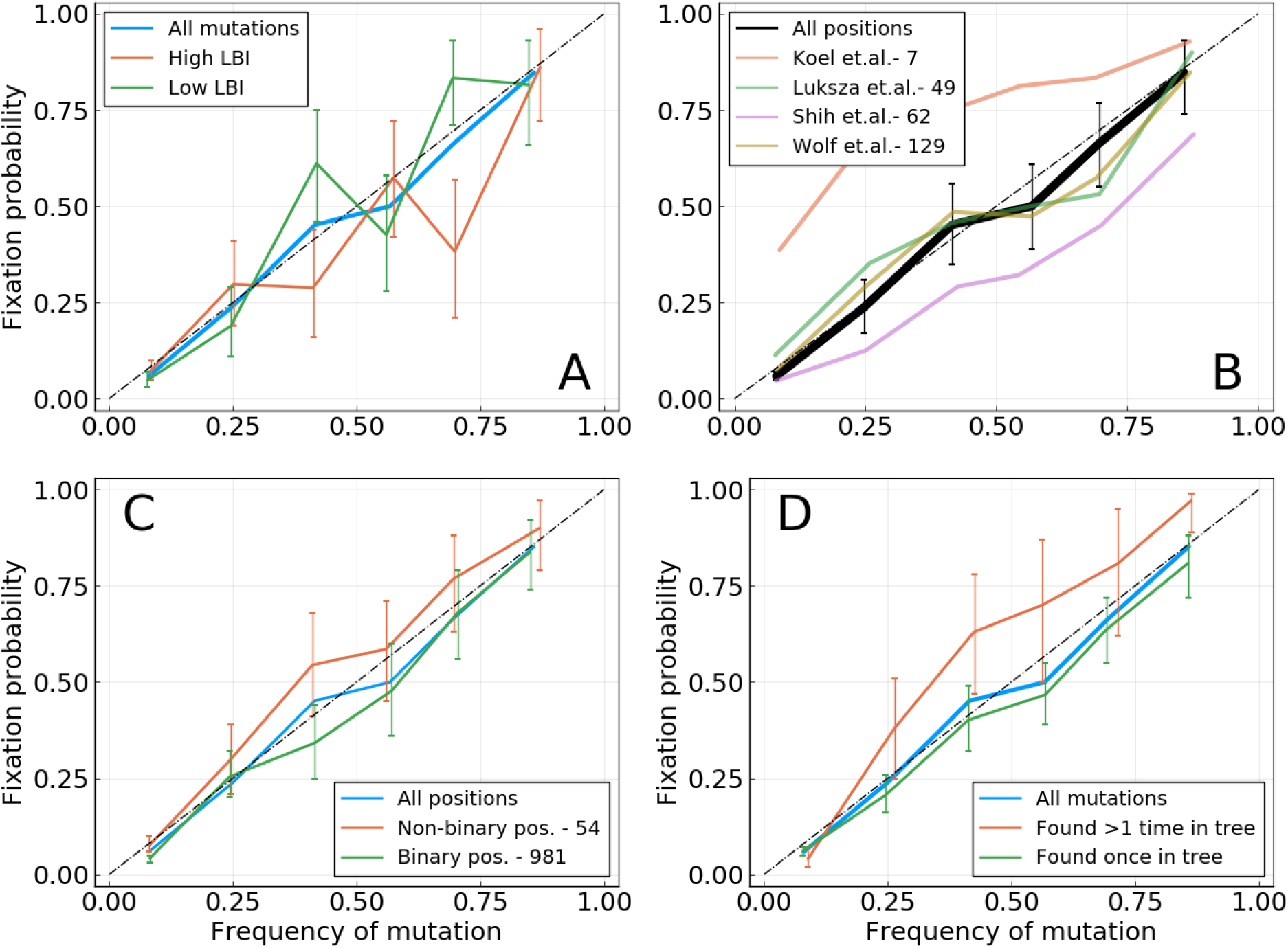
Equivalent to figure 3 of the main text, but with strains carrying the HA1 159S mutation removed.

### 5. Probability of fixation in single locus model of evolution

In [18], Kimura investigates a simple model of evolution with a single locus and a population of size *N*. In this framework, a mutation at this locus with fitness effect *s* and observed at frequency *f* has the following probability of fixation:

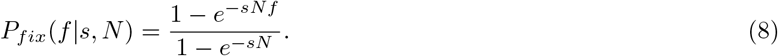

Expanding this formula for *sN «* 1, that is in the weak selection regime, yields at the first order

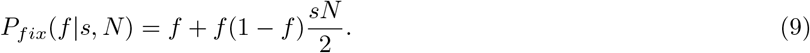

Equation 9 tells us two things. First, when the mutation is neutral, that is *s* = 0, we have *P_fix_*(*f*) = *f*. This naturally confirms the result obtained for a neutral model of evolution. Seconds, when *sN* ≠ 0, we can expect deviations from the diagonal in a *P_fix_* against *f* plot. The sign of these deviations is determined by the sign of *s*, with beneficial mutations being found above diagonal while deleterious one are found below. The amplitude of these deviations depends on the strength of selection *sN*, as well as on the frequency through the *f* (1 *− f*) term, making them larger for *f ∼* 0.5.

### 6. Mutation tables

**Table. S I.**
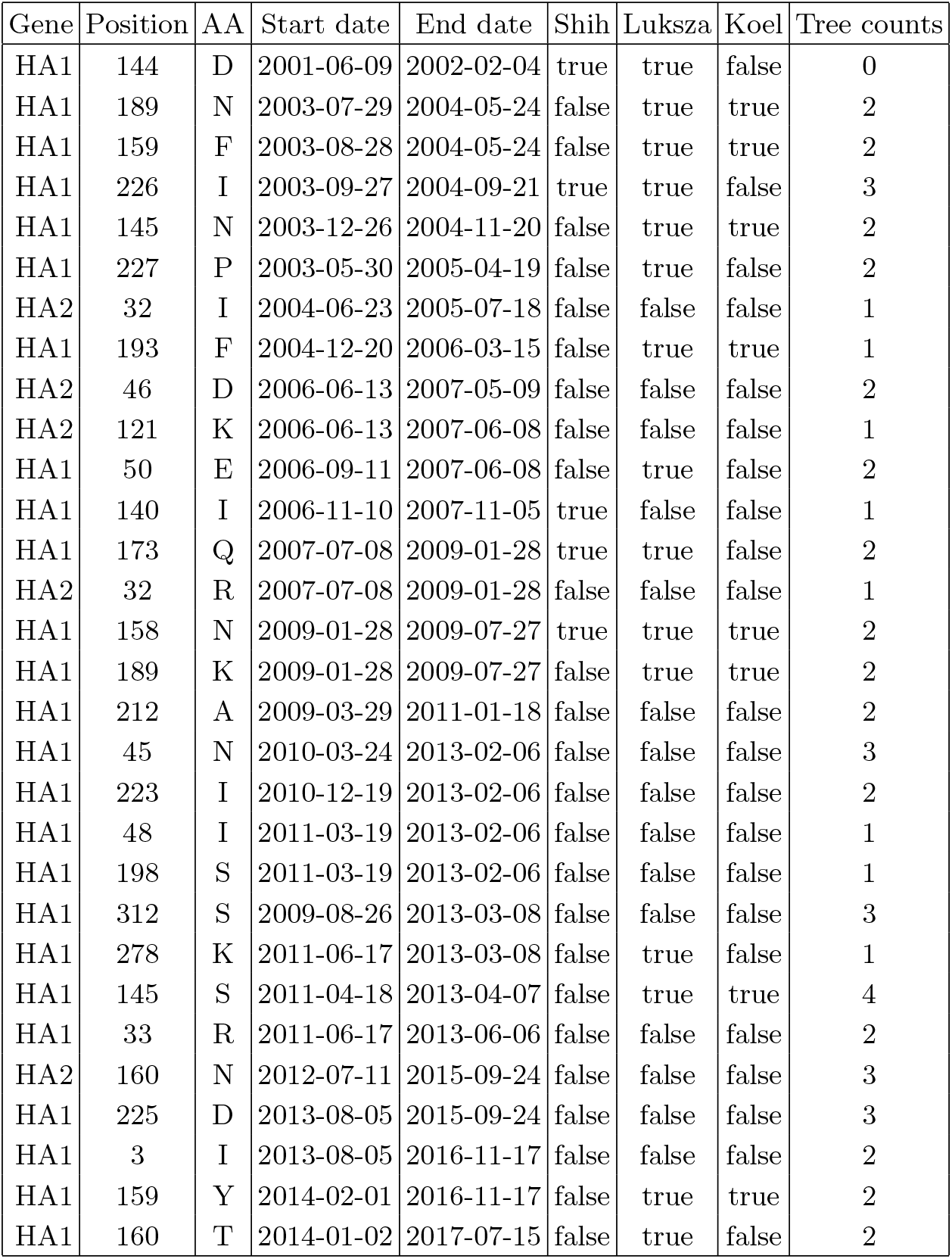
The 30 trajectories that took place between year 2000 and year 2018 and resulted in fixation. Columns `Shih, Luksza` and `Koel` respectively indicate whether the position is found in the epitopes lists in (respectively) [3], [11] and [5]. The `Tree counts` column indicates the number of times the mutation corresponding to the trajectory can be found in the phylogenetic tree. Note that a trajectory is only shown in the table if the sequenced population counts more than 10 strains at its time of fixation. This explains that only 30 trajectories are displayed, whereas more mutations did fix in this period of time.

| Gene | Position | AA | Start date | End date | Shih | Luksza | Koel | Tree counts |
| --- | --- | --- | --- | --- | --- | --- | --- | --- |
| HA1 | 144 | D | 2001-06-09 | 2002-02-04 | true | true | false | 0 |
| HA1 | 189 | N | 2003-07-29 | 2004-05-24 | false | true | true | 2 |
| HA1 | 159 | F | 2003-08-28 | 2004-05-24 | false | true | true | 2 |
| HA1 | 226 | I | 2003-09-27 | 2004-09-21 | true | true | false | 3 |
| HA1 | 145 | N | 2003-12-26 | 2004-11-20 | false | true | true | 2 |
| HA1 | 227 | P | 2003-05-30 | 2005-04-19 | false | true | false | 2 |
| HA2 | 32 | I | 2004-06-23 | 2005-07-18 | false | false | false | 1 |
| HA1 | 193 | F | 2004-12-20 | 2006-03-15 | false | true | true | 1 |
| HA2 | 46 | D | 2006-06-13 | 2007-05-09 | false | false | false | 2 |
| HA2 | 121 | K | 2006-06-13 | 2007-06-08 | false | false | false | 1 |
| HA1 | 50 | E | 2006-09-11 | 2007-06-08 | false | true | false | 2 |
| HA1 | 140 | I | 2006-11-10 | 2007-11-05 | true | false | false | 1 |
| HA1 | 173 | Q | 2007-07-08 | 2009-01-28 | true | true | false | 2 |
| HA2 | 32 | R | 2007-07-08 | 2009-01-28 | false | false | false | 1 |
| HA1 | 158 | N | 2009-01-28 | 2009-07-27 | true | true | true | 2 |
| HA1 | 189 | K | 2009-01-28 | 2009-07-27 | false | true | true | 2 |
| HA1 | 212 | A | 2009-03-29 | 2011-01-18 | false | false | false | 2 |
| HA1 | 45 | N | 2010-03-24 | 2013-02-06 | false | false | false | 3 |
| HA1 | 223 | I | 2010-12-19 | 2013-02-06 | false | false | false | 2 |
| HA1 | 48 | I | 2011-03-19 | 2013-02-06 | false | false | false | 1 |
| HA1 | 198 | S | 2011-03-19 | 2013-02-06 | false | false | false | 1 |
| HA1 | 312 | S | 2009-08-26 | 2013-03-08 | false | false | false | 3 |
| HA1 | 278 | K | 2011-06-17 | 2013-03-08 | false | true | false | 1 |
| HA1 | 145 | S | 2011-04-18 | 2013-04-07 | false | true | true | 4 |
| HA1 | 33 | R | 2011-06-17 | 2013-06-06 | false | false | false | 2 |
| HA2 | 160 | N | 2012-07-11 | 2015-09-24 | false | false | false | 3 |
| HA1 | 225 | D | 2013-08-05 | 2015-09-24 | false | false | false | 3 |
| HA1 | 3 | I | 2013-08-05 | 2016-11-17 | false | false | false | 2 |
| HA1 | 159 | Y | 2014-02-01 | 2016-11-17 | false | true | true | 2 |
| HA1 | 160 | T | 2014-01-02 | 2017-07-15 | false | true | false | 2 |

**Table. S II.** Trajectories of mutations at epitope positions in [3] (*Shih et. al.*) that have been observed at least once above frequency 0.25. The `Fixation` column indicates whether the mutation has fixed, disappeared, or is still polymorphic as of October 2018. The `Max.freq.` column indicates the maximum frequency reached by the trajectory. A maximum frequency of 1 for mutations that finally disappear is explained by trajectories reaching frequency 1 for one time bin and going back to lower values for following ones (a frequency above 0.95 for two time bins in a row defines fixation).

| Gene | Position | AA | Start date | End date | Fixation | Max. freq. |
| --- | --- | --- | --- | --- | --- | --- |
| HA1 | 106 | A | 2001-02-09 | 2002-02-04 | lost | 1.0 |
| HA1 | 144 | D | 2001-06-09 | 2002-02-04 | fixed | 1.0 |
| HA1 | 105 | H | 2003-04-30 | 2003-10-27 | lost | 1.0 |
| HA1 | 126 | D | 2003-04-30 | 2004-05-24 | lost | 1.0 |
| HA1 | 140 | Q | 2004-01-25 | 2004-06-23 | lost | 0.31 |
| HA1 | 226 | I | 2003-09-27 | 2004-09-21 | fixed | 1.0 |
| HA1 | 173 | E | 2004-12-20 | 2006-03-15 | lost | 0.63 |
| HA1 | 142 | G | 2006-06-13 | 2007-05-09 | lost | 0.71 |
| HA1 | 144 | D | 2006-07-13 | 2007-05-09 | lost | 0.67 |
| HA1 | 128 | A | 2006-09-11 | 2007-05-09 | lost | 0.25 |
| HA1 | 157 | S | 2006-09-11 | 2007-05-09 | lost | 0.59 |
| HA1 | 140 | I | 2006-11-10 | 2007-11-05 | fixed | 1.0 |
| HA1 | 173 | N | 2007-12-05 | 2008-07-02 | lost | 0.3 |
| HA1 | 157 | S | 2007-12-05 | 2008-09-30 | lost | 0.31 |
| HA1 | 173 | E | 2006-06-13 | 2008-12-29 | lost | 0.67 |
| HA1 | 173 | Q | 2007-07-08 | 2009-01-28 | fixed | 0.96 |
| HA1 | 158 | N | 2009-01-28 | 2009-07-27 | fixed | 0.96 |
| HA1 | 62 | K | 2009-01-28 | 2011-05-18 | lost | 0.73 |
| HA1 | 144 | K | 2009-01-28 | 2011-05-18 | lost | 0.75 |
| HA1 | 62 | V | 2011-04-18 | 2011-09-15 | lost | 0.34 |
| HA1 | 157 | S | 2013-05-07 | 2015-09-24 | lost | 0.35 |
| HA1 | 128 | A | 2012-08-10 | 2016-11-17 | lost | 0.81 |
| HA1 | 197 | K | 2015-11-23 | 2016-11-17 | lost | 0.27 |
| HA1 | 142 | R | 2018-05-11 | 2018-10-08 | lost | 0.38 |
| HA1 | 142 | G | 2012-03-13 |  | poly | 0.86 |
| HA1 | 144 | S | 2013-12-03 |  | poly | 0.96 |
| HA1 | 121 | K | 2015-12-23 |  | poly | 0.82 |
| HA1 | 142 | K | 2016-05-21 |  | poly | 0.77 |
| HA1 | 62 | G | 2017-03-17 |  | poly | 0.75 |
| HA1 | 128 | A | 2018-01-11 |  | poly | 0.56 |

**Table. S III.** Same as table SII, for [11] (*Luksza et. al.*).

| Gene | Position | AA | Start date | End date | Fixation | Max. freq. |
| --- | --- | --- | --- | --- | --- | --- |
| HA1 | 50 | G | 2001-02-09 | 2002-02-04 | lost | 1.0 |
| HA1 | 144 | D | 2001-06-09 | 2002-02-04 | fixed | 1.0 |
| HA1 | 126 | D | 2003-04-30 | 2004-05-24 | lost | 1.0 |
| HA1 | 189 | N | 2003-07-29 | 2004-05-24 | fixed | 1.0 |
| HA1 | 159 | F | 2003-08-28 | 2004-05-24 | fixed | 1.0 |
| HA1 | 226 | I | 2003-09-27 | 2004-09-21 | fixed | 1.0 |
| HA1 | 145 | N | 2003-12-26 | 2004-11-20 | fixed | 1.0 |
| HA1 | 188 | N | 2004-07-23 | 2005-02-18 | lost | 0.36 |
| HA1 | 227 | P | 2003-05-30 | 2005-04-19 | fixed | 1.0 |
| HA1 | 173 | E | 2004-12-20 | 2006-03-15 | lost | 0.63 |
| HA1 | 193 | F | 2004-12-20 | 2006-03-15 | fixed | 0.97 |
| HA1 | 142 | G | 2006-06-13 | 2007-05-09 | lost | 0.71 |
| HA1 | 144 | D | 2006-07-13 | 2007-05-09 | lost | 0.67 |
| HA1 | 157 | S | 2006-09-11 | 2007-05-09 | lost | 0.59 |
| HA1 | 50 | E | 2006-09-11 | 2007-06-08 | fixed | 0.95 |
| HA1 | 173 | N | 2007-12-05 | 2008-07-02 | lost | 0.3 |
| HA1 | 157 | S | 2007-12-05 | 2008-09-30 | lost | 0.31 |
| HA1 | 173 | E | 2006-06-13 | 2008-12-29 | lost | 0.67 |
| HA1 | 173 | Q | 2007-07-08 | 2009-01-28 | fixed | 0.96 |
| HA1 | 158 | N | 2009-01-28 | 2009-07-27 | fixed | 0.96 |
| HA1 | 189 | K | 2009-01-28 | 2009-07-27 | fixed | 0.96 |
| HA1 | 213 | A | 2009-01-28 | 2010-02-22 | lost | 0.68 |
| HA1 | 144 | K | 2009-01-28 | 2011-05-18 | lost | 0.75 |
| HA1 | 53 | N | 2009-11-24 | 2013-02-06 | lost | 0.72 |
| HA1 | 278 | K | 2011-06-17 | 2013-03-08 | fixed | 0.98 |
| HA1 | 145 | S | 2011-04-18 | 2013-04-07 | fixed | 0.99 |
| HA1 | 159 | S | 2013-11-03 | 2015-08-25 | lost | 0.46 |
| HA1 | 157 | S | 2013-05-07 | 2015-09-24 | lost | 0.35 |
| HA1 | 159 | Y | 2014-02-01 | 2016-11-17 | fixed | 0.97 |
| HA1 | 159 | S | 2015-10-24 | 2016-11-17 | lost | 0.4 |
| HA1 | 197 | K | 2015-11-23 | 2016-11-17 | lost | 0.27 |
| HA1 | 160 | T | 2014-01-02 | 2017-07-15 | fixed | 0.96 |
| HA1 | 142 | R | 2018-05-11 | 2018-10-08 | lost | 0.38 |
| HA1 | 135 | N | 2018-06-10 | 2018-10-08 | lost | 0.38 |
| HA1 | 142 | G | 2012-03-13 |  | poly | 0.86 |
| HA1 | 144 | S | 2013-12-03 |  | poly | 0.96 |
| HA1 | 121 | K | 2015-12-23 |  | poly | 0.82 |
| HA1 | 142 | K | 2016-05-21 |  | poly | 0.77 |
| HA1 | 131 | K | 2016-09-18 |  | poly | 0.77 |
| HA1 | 135 | K | 2016-11-17 |  | poly | 0.47 |

**Table. S IV.**
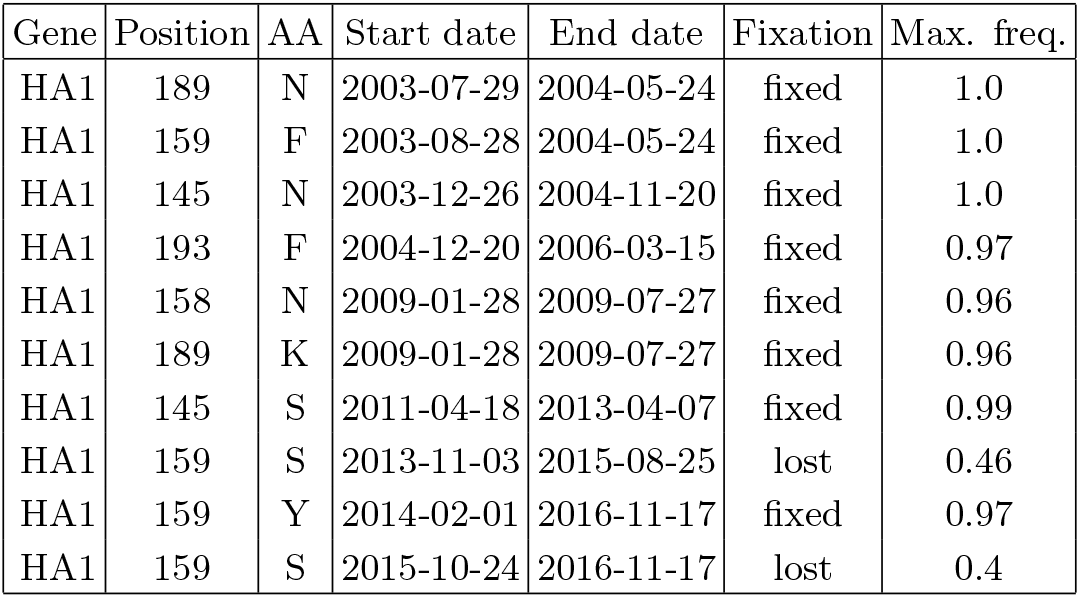
Same as table SII, for [5] (*Koel et. al.*).

### 7. Supplementary figures

**Figure S5.**
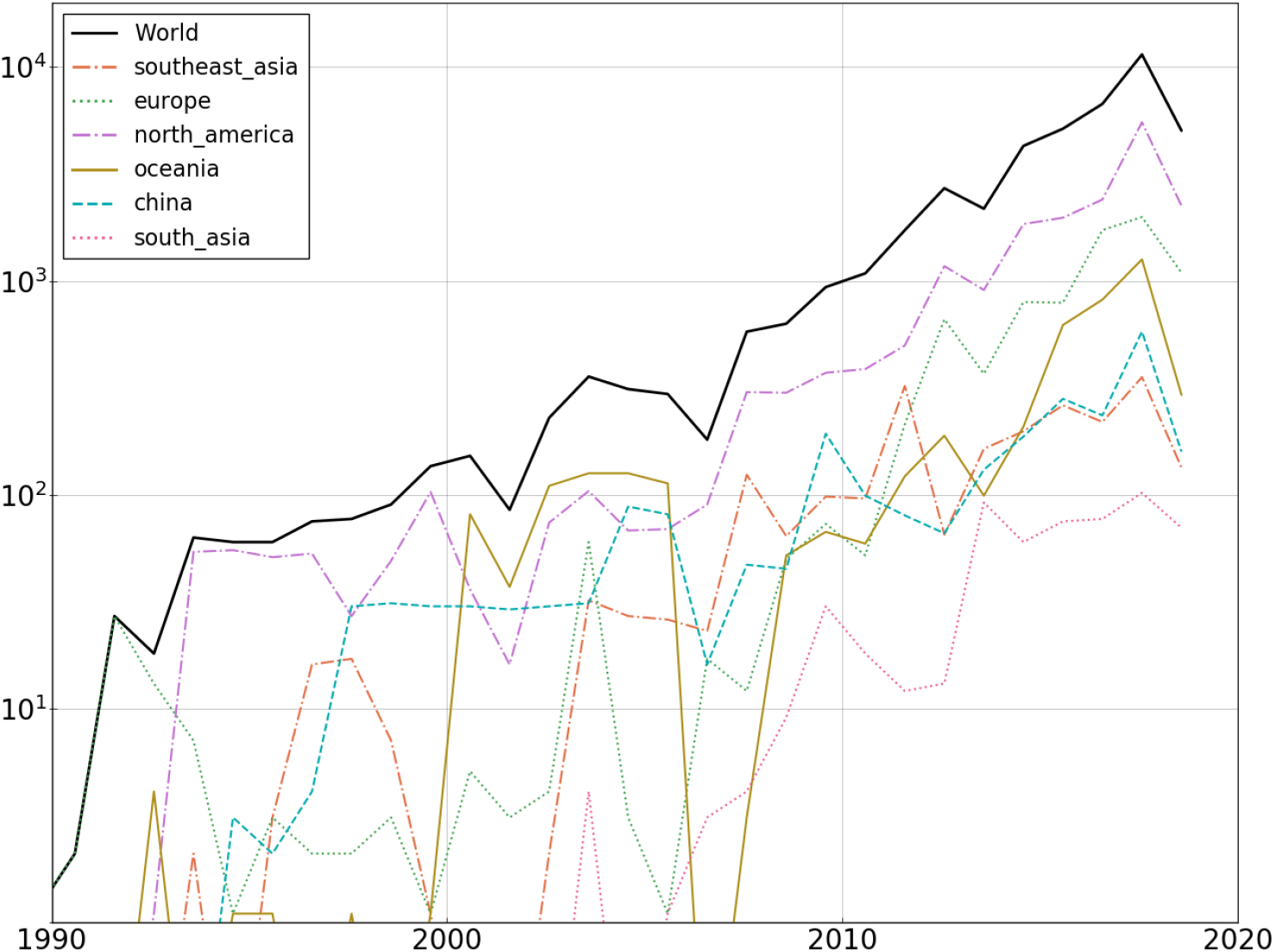
Number of A/H3N2 HA sequences per year from year 1990.

**Figure S6.**
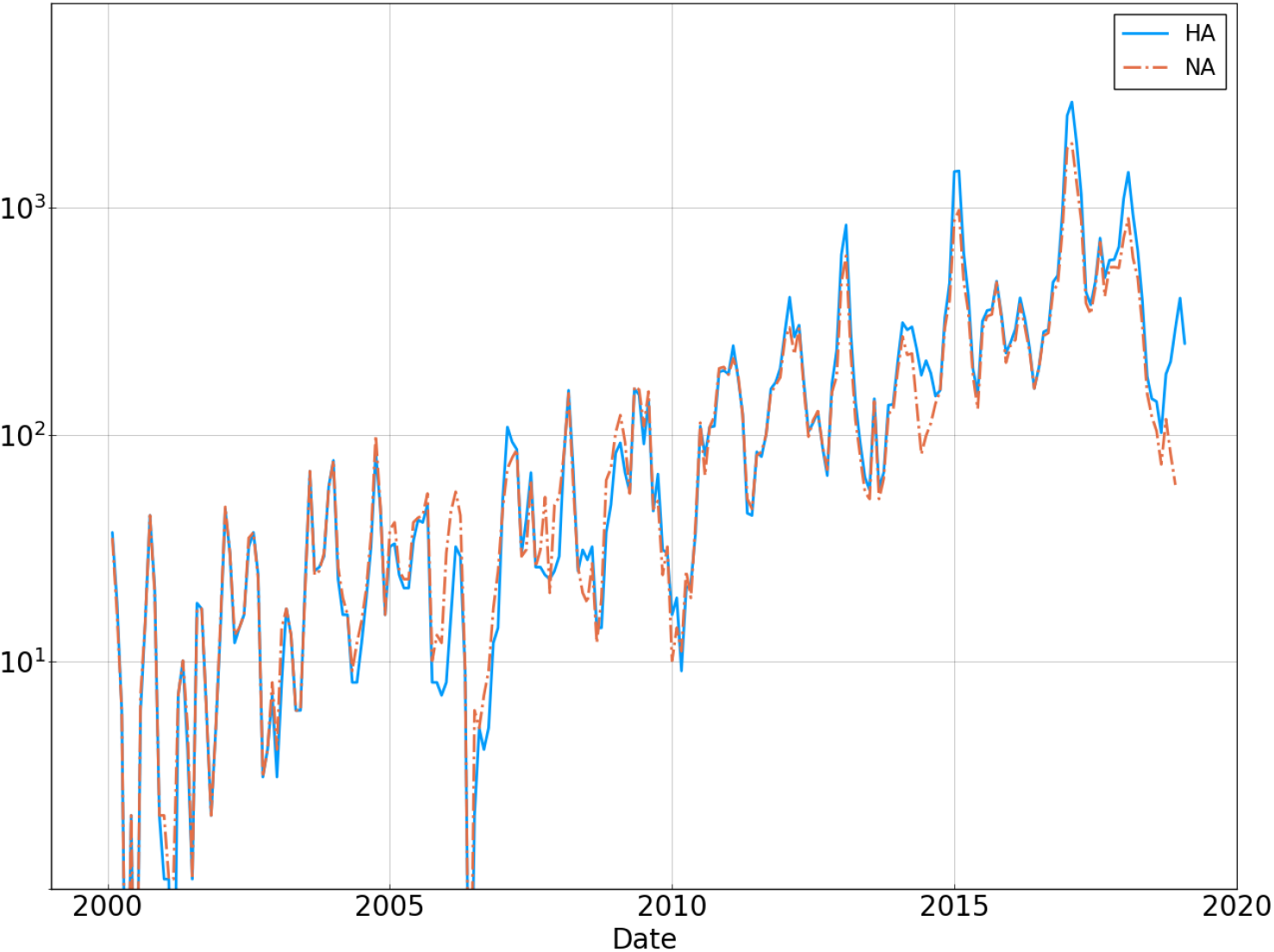
Number of H3N2 HA and NA sequences per month from year 2000.

**Figure S7.**
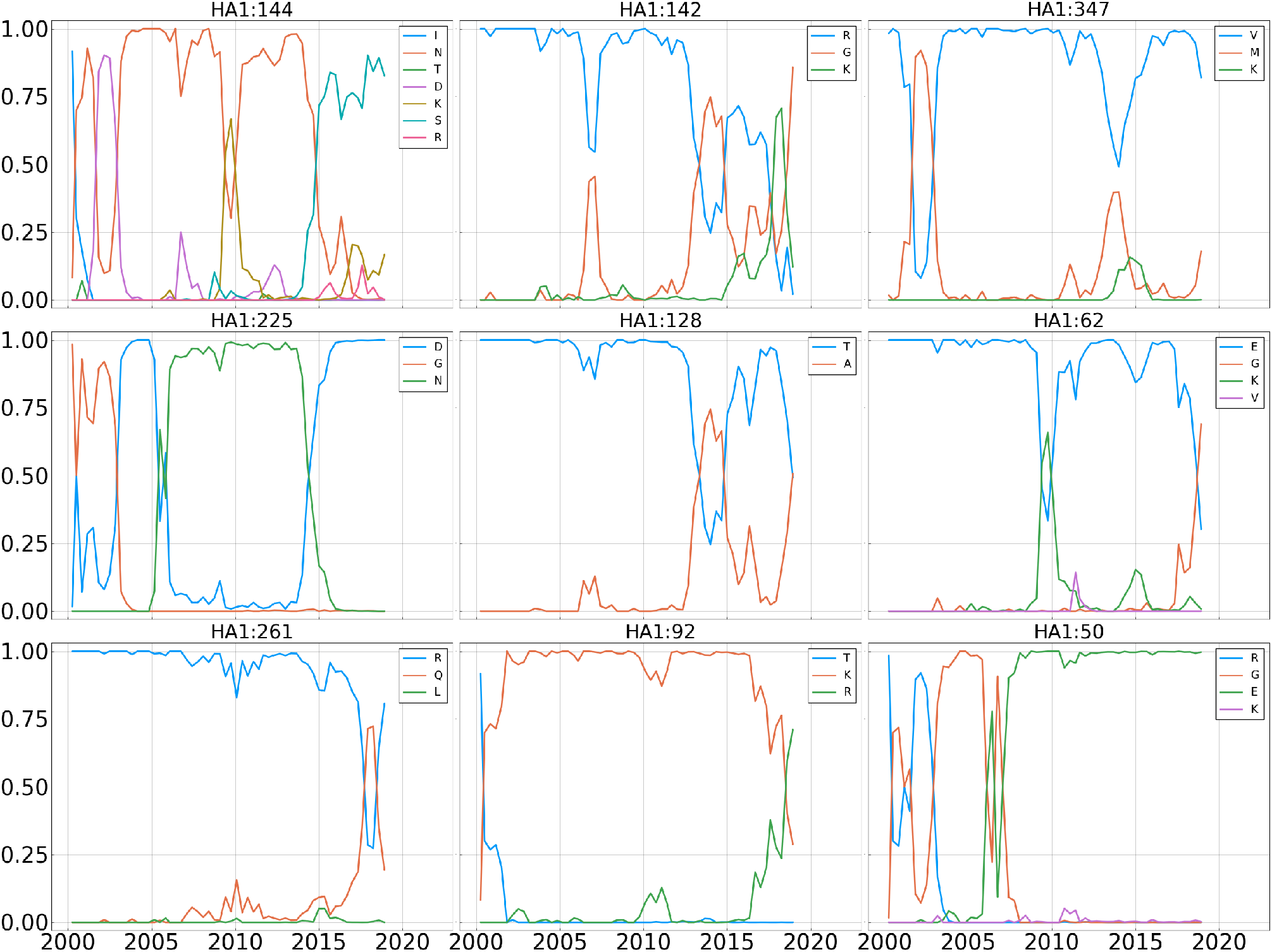
Frequency trajectories for the 9 most entropic positions in the A/H3N2 HA protein.

**Figure S8.**
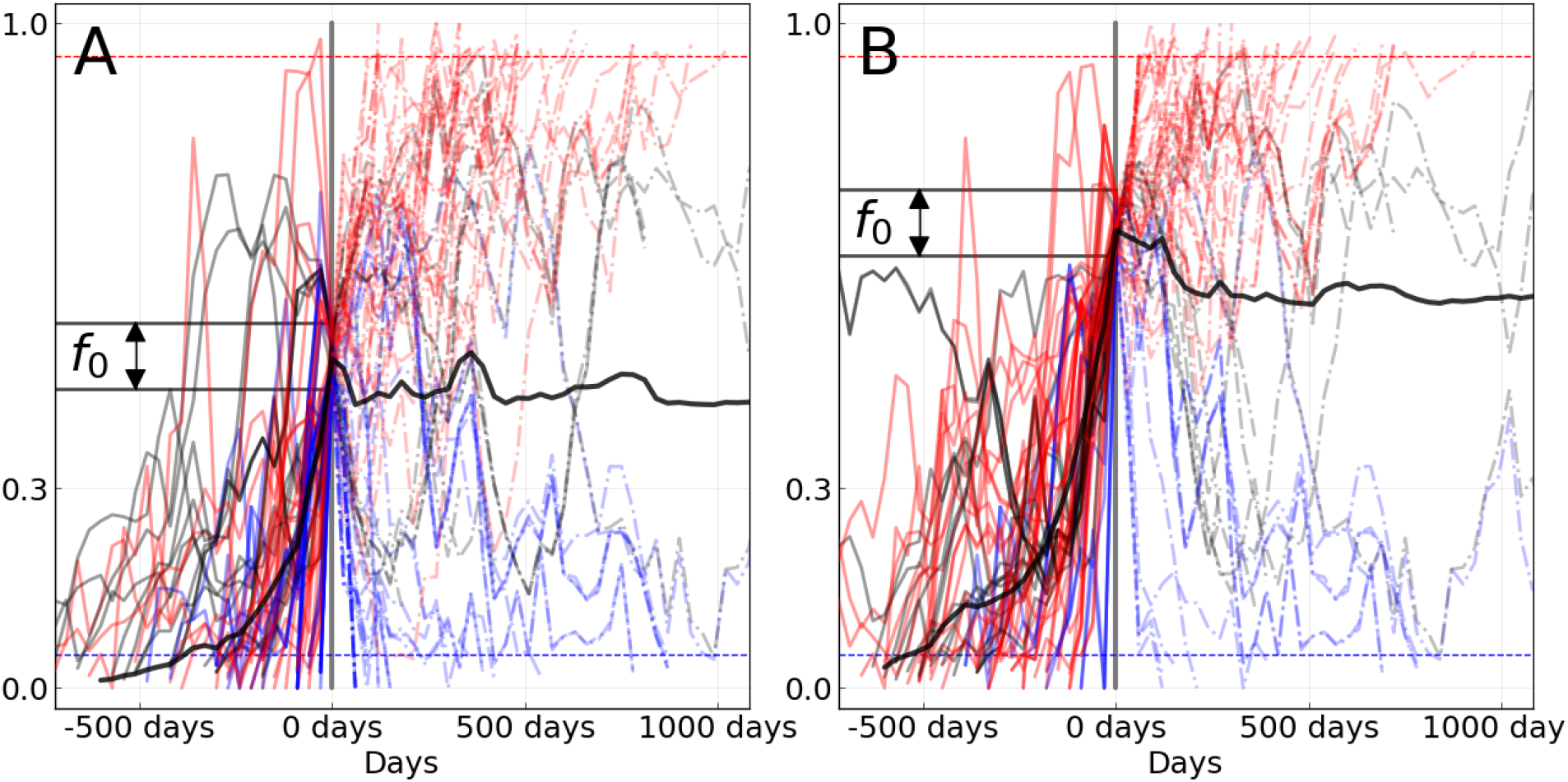
Equivalent to panel **B** of figure 1 of the main text for A/H3N2, with *f*_0_ equal 0.5 in **A**(76 trajectories), and 0.7 in **B**(63 trajectories).

**Figure S9.**
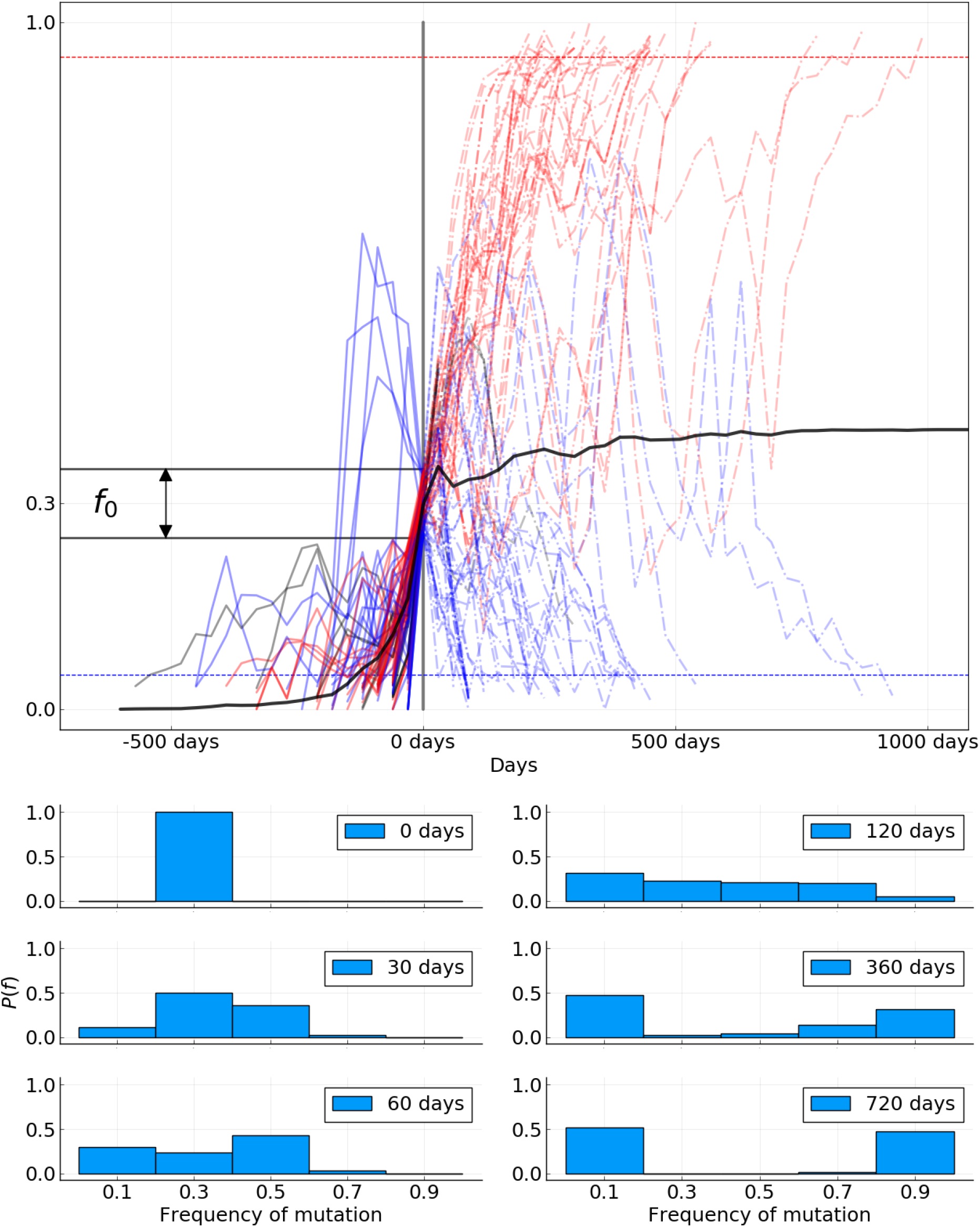
Equivalent to panels **B** and **C** of figure 1 of the main text for A/H1N1pdm influenza. 89 trajectories are shown and participate to the mean (thick black line).

**Figure S10.**
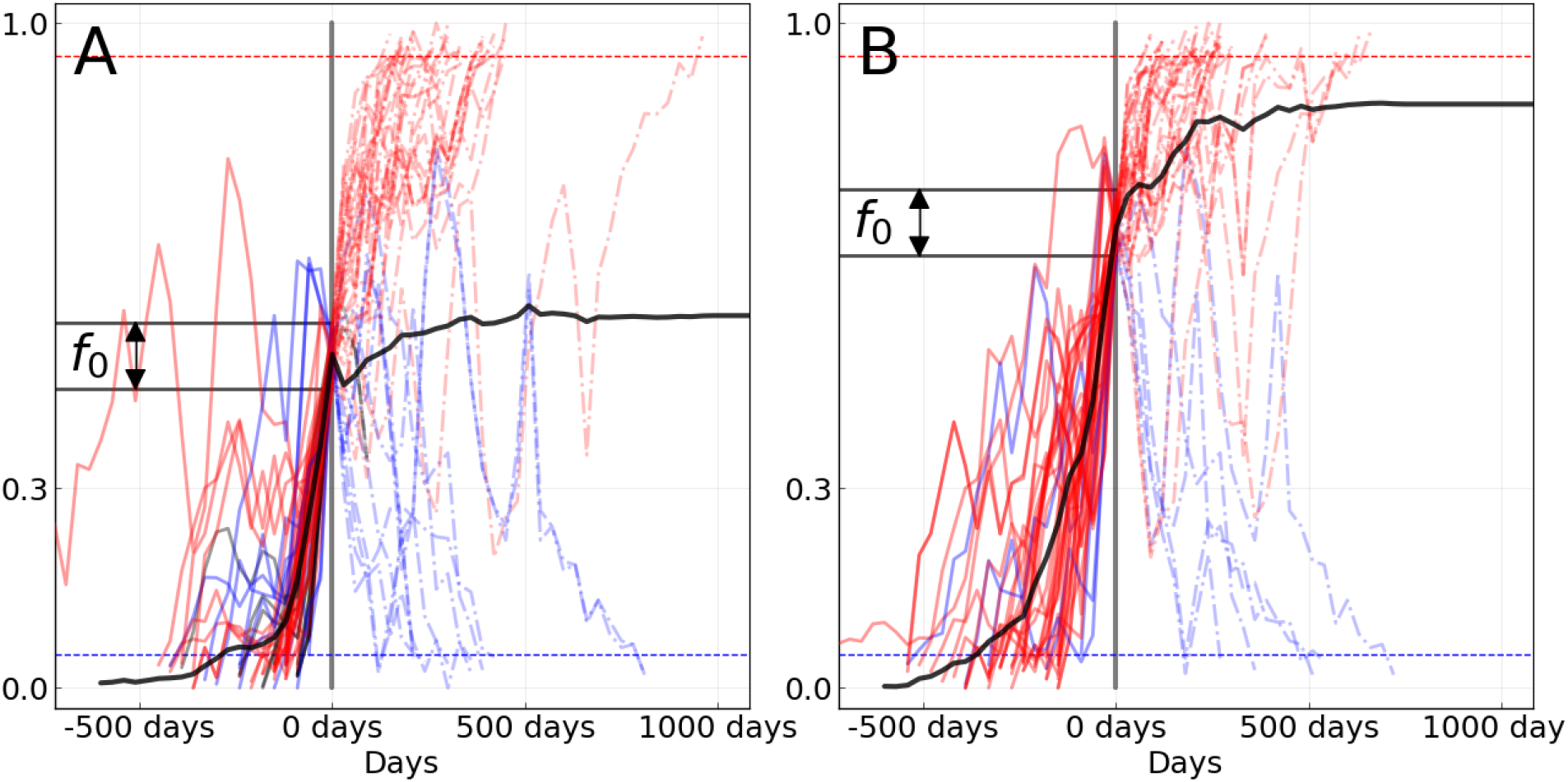
Equivalent to panel **B** of figure 1 of the main text for A/H1N1pdm, with *f*_0_ equal 0.5 in **A**(50 trajectories), and 0.7 in **B**(41 trajectories).

**Figure S11.**
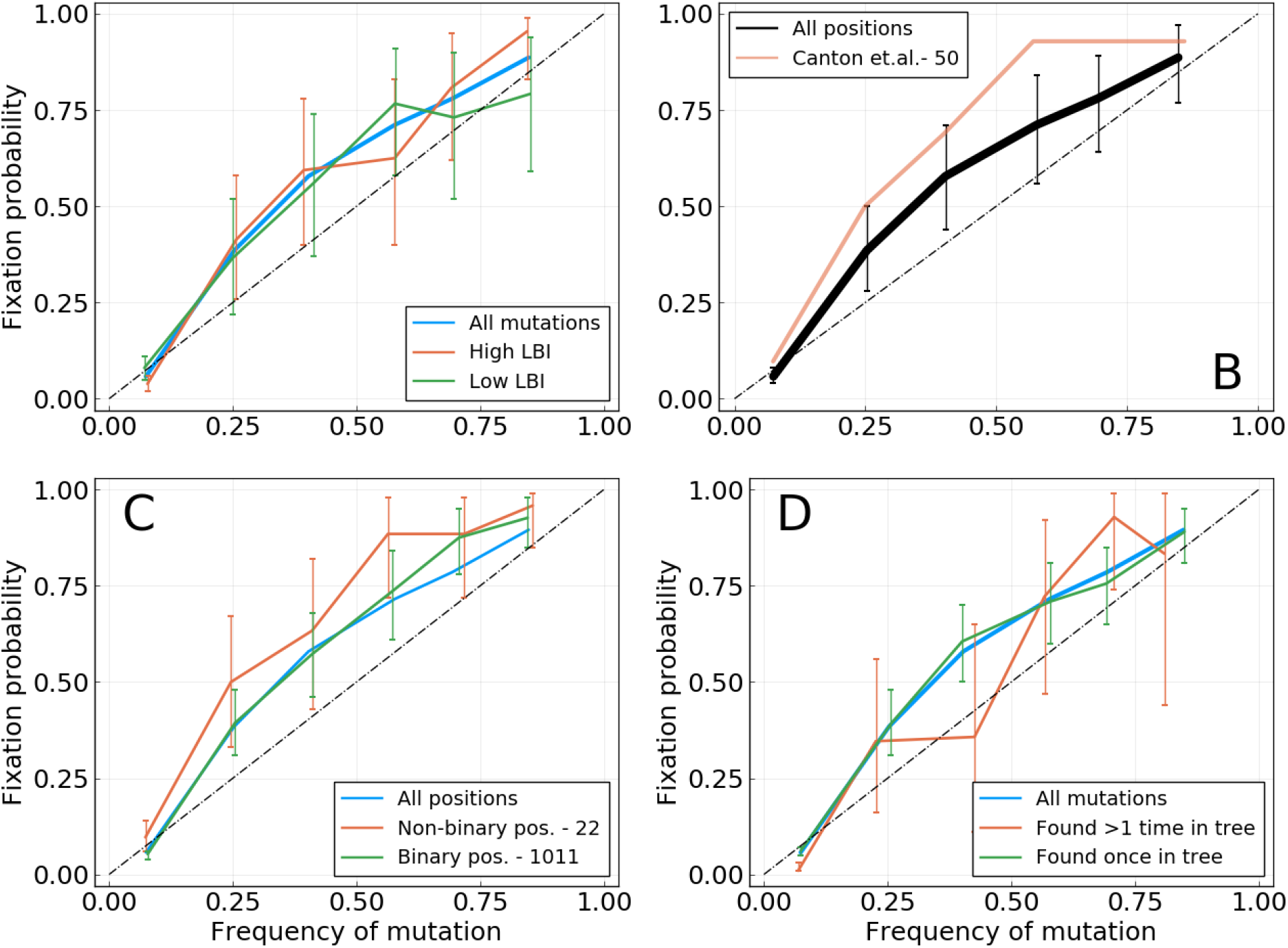
Equivalent of figure 3 of the main text for the HA gene of A/H1N1pdm influenza. Fixation probability *P_fix_*(*f*) as a function of frequency. **A**: Mutation with higher or lower LBI values, based on their position with respect to the median LBI value. **B**: Different lists of epitope positions in the HA protein. The authors and the number of positions is indicated in the legend. **C**: Mutations for binary positions, *i.e.* positions for which we never see more than two amino acids in the same time bin. **D**: Mutations that appear once or more than once in the tree for a given time bin.

**Figure S12.**
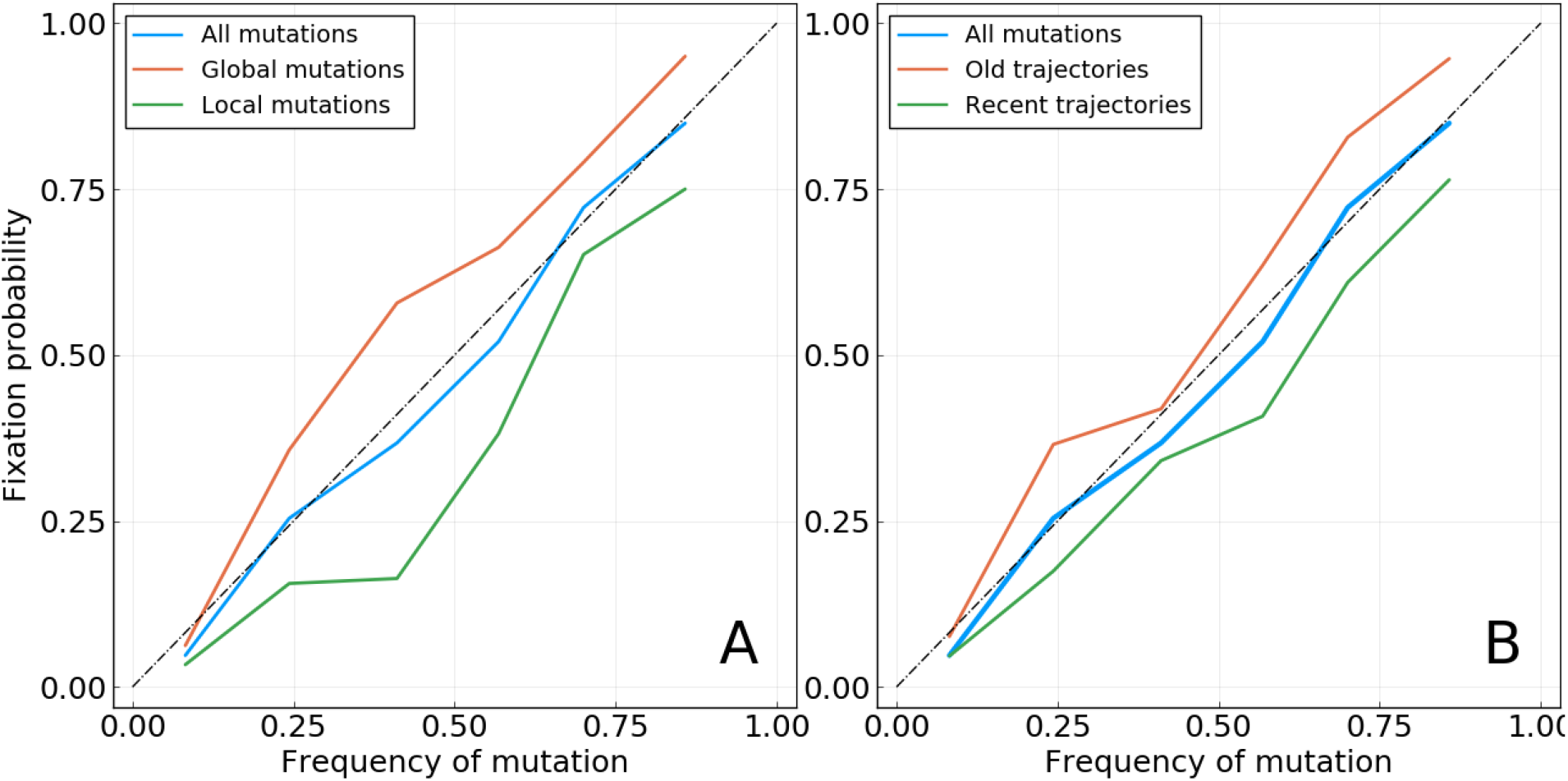
Based on A/H3N2 HA and NA. **A**: Mutations with a higher or lower geographical spread, based on the median value of the score used (see Methods). *Note*: the words *local* and *global* only reflect the position of the geographic spread of the mutation relative to the median value computed for all mutations found at this frequency. As this median value may change with the considered frequency bin, so does the definition of local and global mutations. **B**: Mutations whose trajectories are older or more recent, based on the median age of trajectories when reaching the considered frequency *f*.

**Figure S13.**
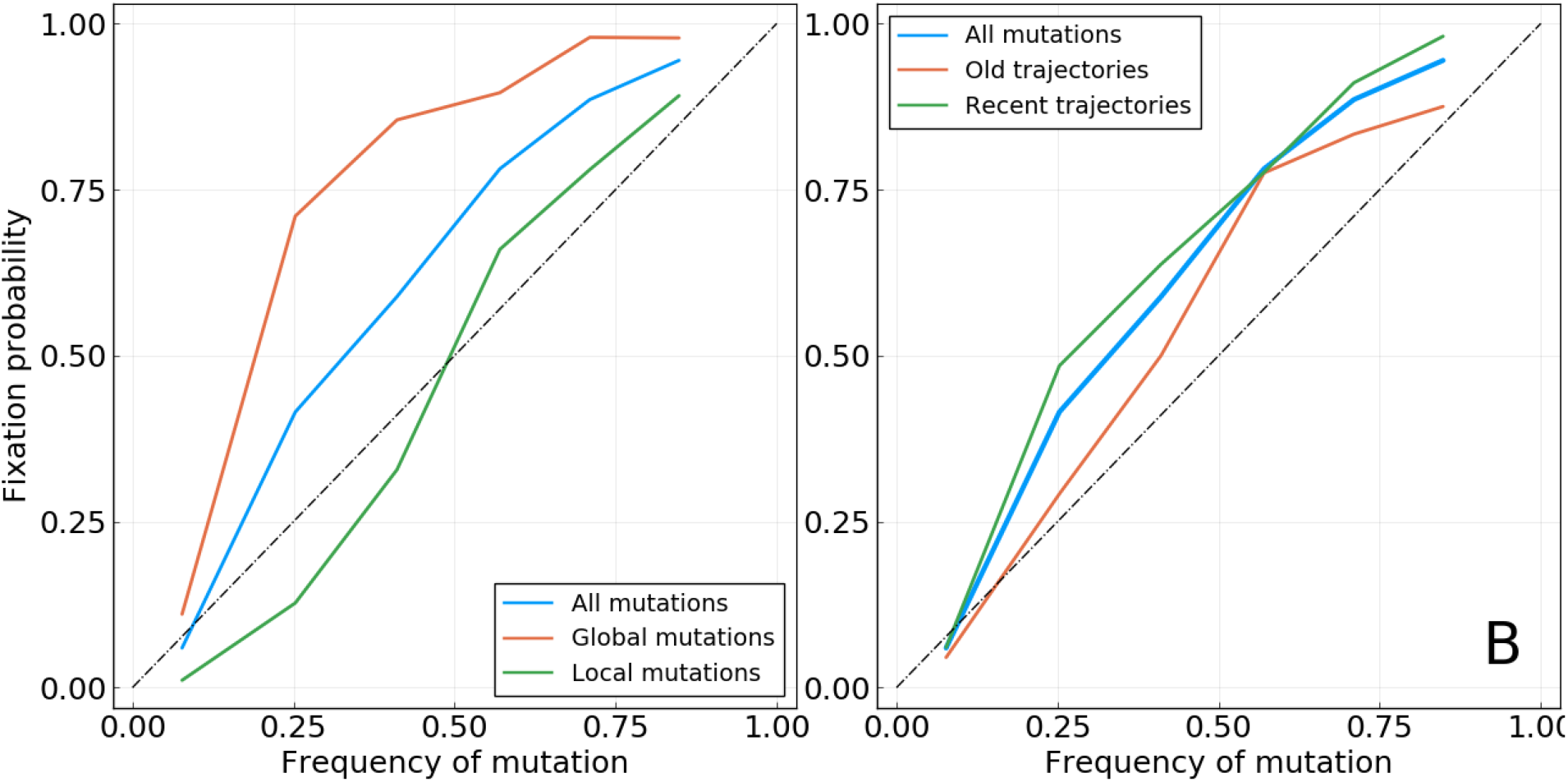
Based on A/H1N1pdm HA and NA. **A**: Mutations with a higher or lower geographical spread, based on the median value of the score used (see Methods). *Note*: the words *local* and *global* only reflect the position of the geographic spread of the mutation relative to the median value computed for all mutations found at this frequency. As this median value may change with the considered frequency bin, so does the definition of local and global mutations. **B**: Mutations whose trajectories are older or more recent, based on the median age of trajectories when reaching the considered frequency *f*.

**Figure S14.**
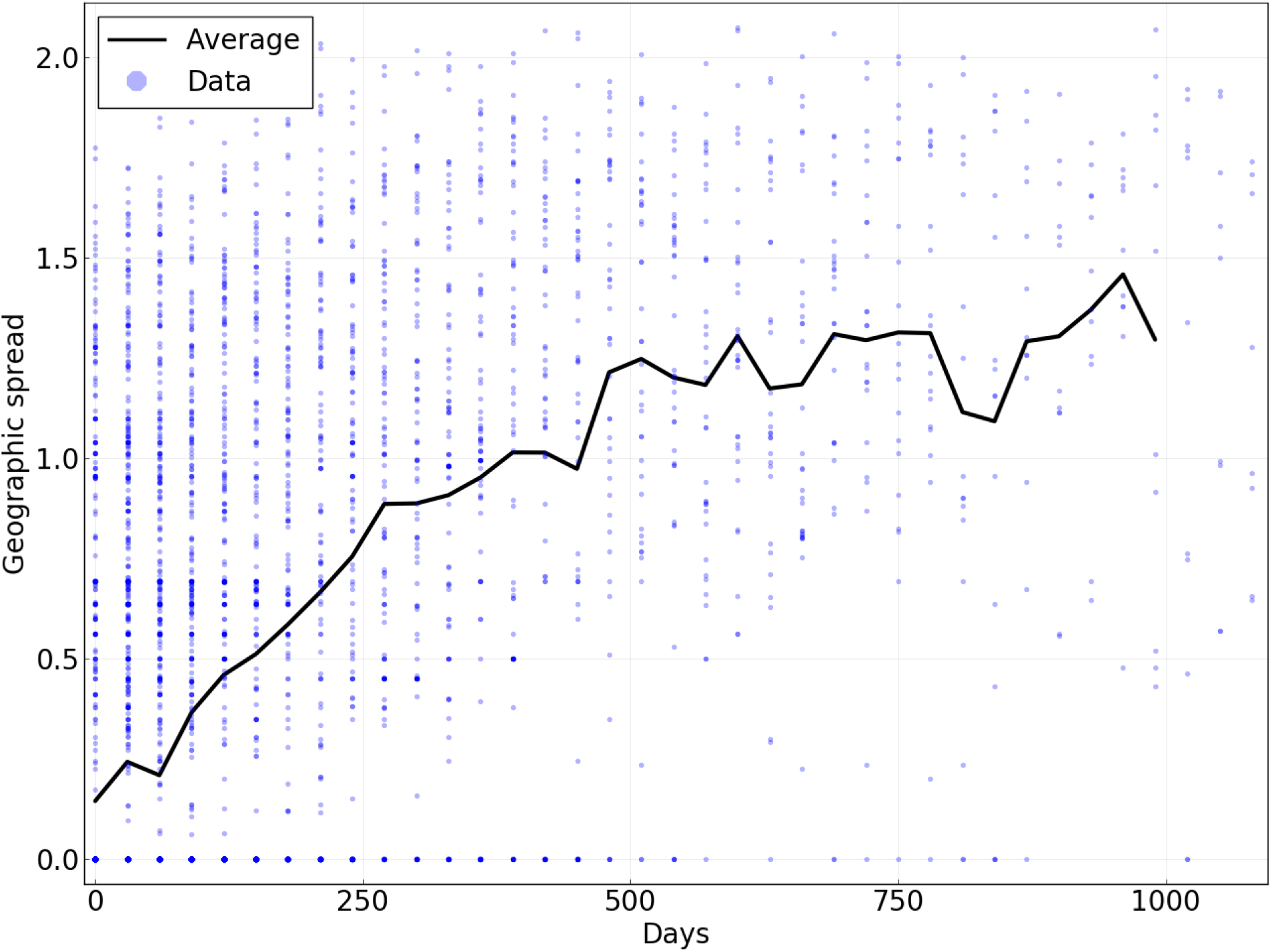
Geographic spread of mutations as a function of the time for which they have been present in the population above a frequency of 5%. Points represent individual mutations and for a population in a given time bin. The line is the average of dots for a given value on the *x*-axis. Based on data for A/H3N2 HA.

**Figure S15.**
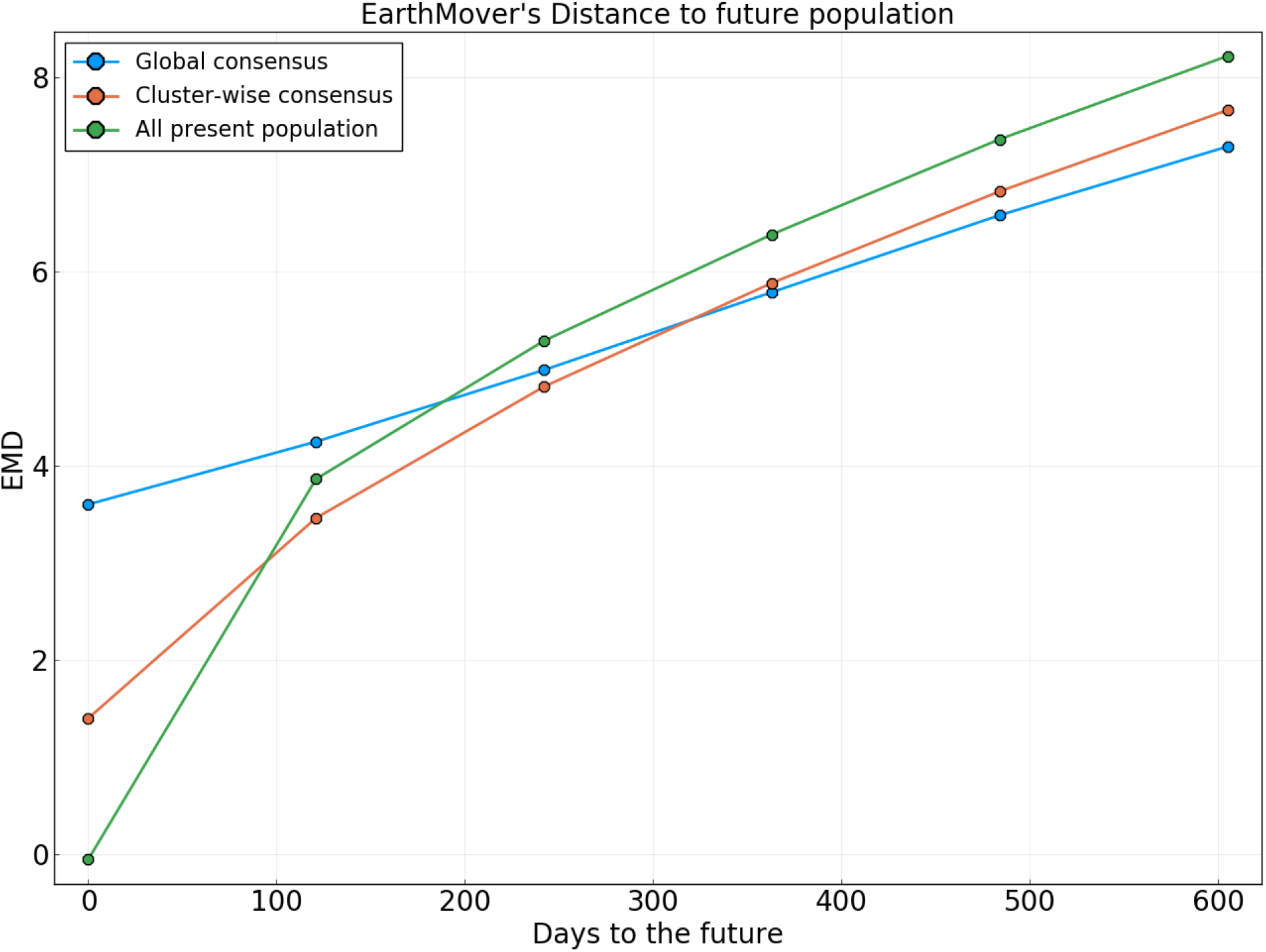
Earth mover’s distance to the future population for different predictors. A present population consists of all A/H3N2 HA sequences sampled in a 4 months time window. Quantities are averaged over all possible “present” populations from the year 2002. Predictors are: **Global consensus**: Consensus sequence of the present population. Best long-term predictor for a structure-less neutrally evolving population. **All present population**: All sequences in the present population. Perfect predictor if the population does not change at all through time. **Cluster-wise consensus**: Consensus sequence for each cluster in the present population. Clusters are based on local maxima of the LBI. Sequences are assigned to a given cluster based on their tree branch-length distance to the corresponding local maximum.

**Figure S16.**
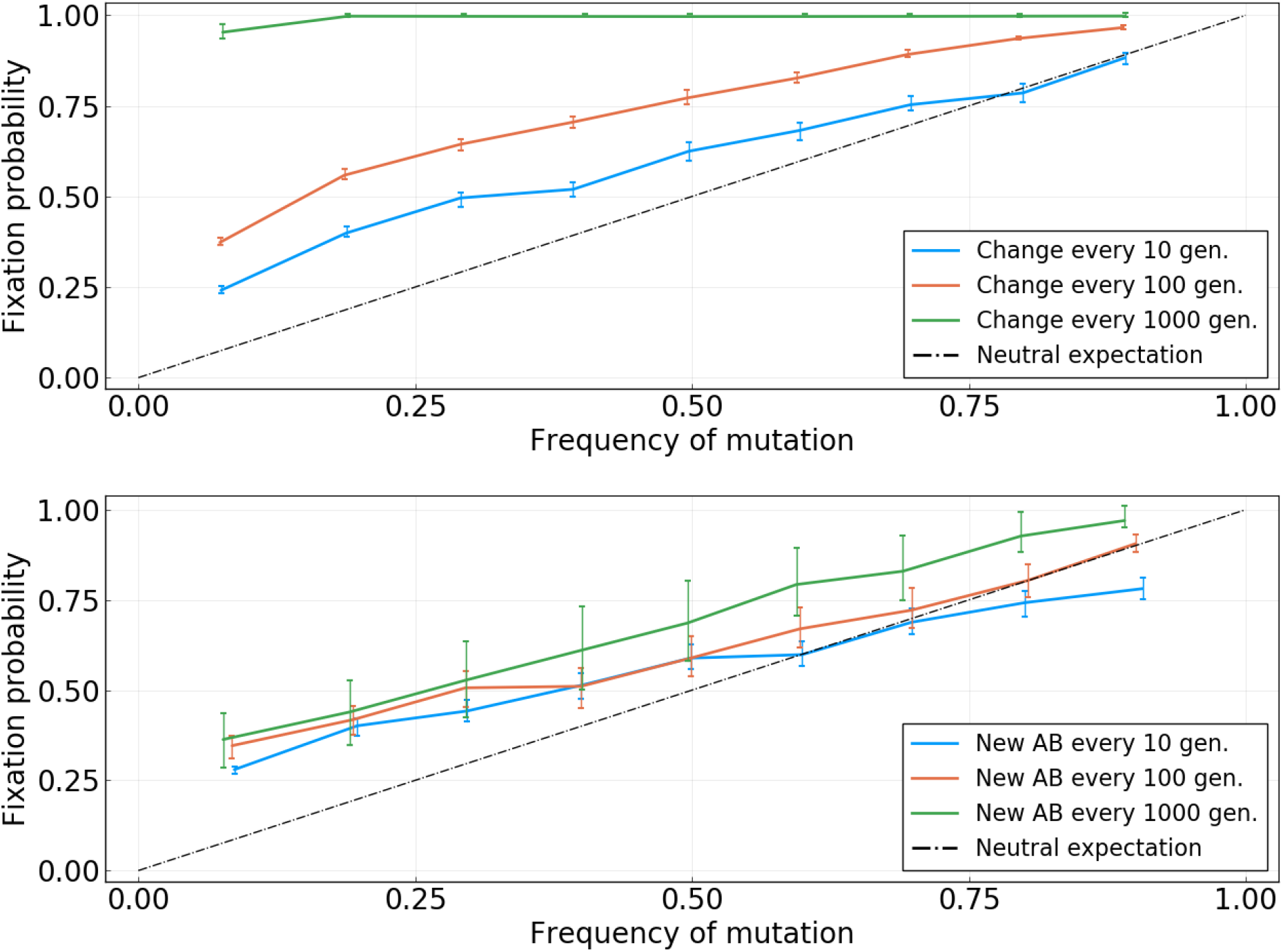
Fixation probability as a function of frequency for the simulations discussed in the main text. **Top:** Simulation without antibodies. The three colored curves reflect different rate of change for the fitness landscape. Visual inspection of the frequency trajectories indicates a typical sweep time of ~400 generations. **Bottom:** Simulation with antibodies. The different colored curves indicate the rate at which antibodies are introduced.

**Figure S17.**
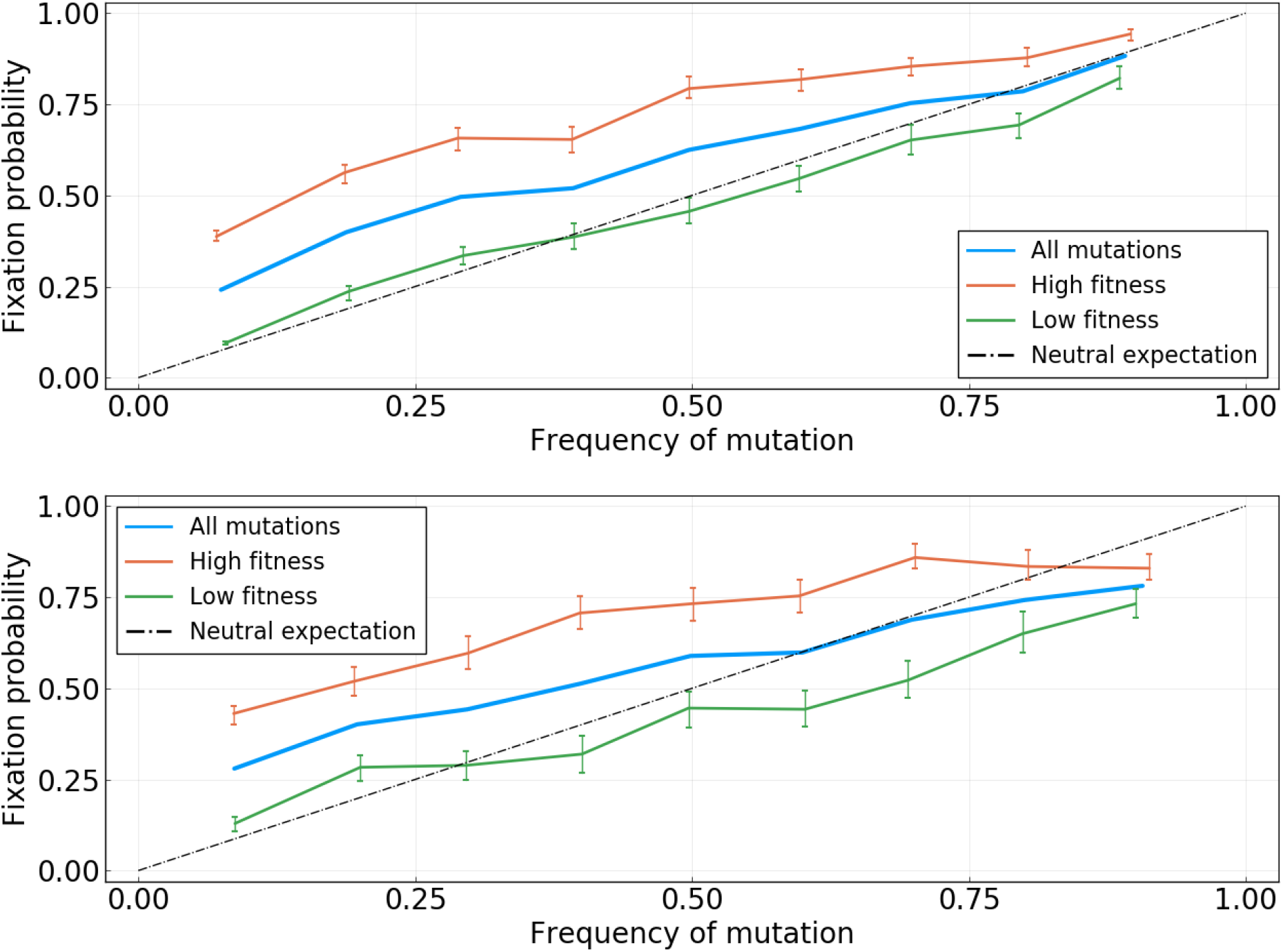
Fixation probability as a function of frequency for the simulations discussed in the main text, with trajectories stratified according to real fitness values. “High” and “low” fitness classes are defined with respect to the median value. **Top:** Simulation without antibodies and with changes to the fitness landscape every *dt* = 10 generations. **Bottom:** Simulation with antibodies, with a new antibody every *dt* = 10 generations.

